# Cysteine Redox State Governs the Condensation Pathway of Hendra Virus W Protein and Differentially Impacts Type I IFN and NF-κB signaling

**DOI:** 10.1101/2024.01.22.576663

**Authors:** Frank Gondelaud, Alexandre Lalande, Giulia Pesce, Carole Meunier, Christophe Bignon, Denis Ptchelkine, Yu Gu, Eva Ogire, Pierre-Yves Lozach, Denis Gerlier, Cyrille Mathieu, Sonia Longhi

## Abstract

The Hendra and Nipah viruses (HeV and NiV) are zoonotic biosafety level-4 pathogens belonging to the *Paramyxoviridae* family. We previously showed that their W protein, a key player in the evasion of the host antiviral response, forms highly flexible, curved fibrils *in vitro*. Here, we show that the cysteine oxidation state acts as a molecular switch controlling the formation of either amorphous aggregates or flexible fibrils, and that residues 2 to 29 are essential for fibrillation. We also uncover that the HeV W protein (W^HeV^) can also self-assemble *in cellula*. W^HeV^ forms distinct types of nuclear condensates that exhibit different dependencies on the cysteine redox-state. While deletion of residues 2-29 prevents formation of nuclear filaments, cysteine-to-serine substitution mainly impairs the formation of non-filamentous condensates. Both infection and W^HeV^ ectopic expression trigger oxidative stress presumably favorable to W^HeV^ condensation. Finally, we show that impaired ability to form redox-sensitive, non-filamentous condensates is associated with a reduced W ability to inhibit the NF-κB pathway, while it conversely enhances W ability to repress the interferon response pathway.

## Introduction

Hendra (HeV) and Nipah viruses (NiV) are members of the *Henipavirus* genus, within the *Paramyxoviridae* family of the *Mononegavirales* order. They are zoonotic viruses responsible for severe encephalitis and respiratory illness with a high case fatality, and as such, they are considered high-priority targets by the WHO. Since the identification of HeV and NiV in humans and fruit-bats from the *Pteropus* genus (Escudero-Pérez et al., 2023; Quarleri et al., 2022), new species have been described including the Cedar virus identified in bats (Marsh et al., 2012), the Langya virus identified in patients, bats and shrews (Amin et al., 2023; Mallapaty, 2022) and two novel species in shrews (Lee et al., 2021). Transmission from bats to humans occurs through the consumption of raw date palm sap (NiV) or from close contact with infected livestock, particularly horses (HeV) and pigs (NiV) (Dawes and Freiberg, 2019). Interhuman transmissions also occurred in the case of NiV Bangladesh strain (Nikolay et al., 2019). Henipavirus outbreaks are unpredictable but occur almost every year with a high case fatality rate (70-100%) while no antiviral therapeutic nor vaccine is currently available (Gómez Román et al., 2022; Quarleri et al., 2022). For these reasons, HeV and NiV are considered as biosafety level-4 pathogens.

Henipaviruses, like all paramyxoviruses, are enveloped viruses containing a non-segmented single-stranded RNA genome of negative polarity (-ssRNA) tightly associated with the nucleoprotein (N). The encapsidated RNA is the template of the viral RNA-dependent RNA polymerase (L) that, in association with the phosphoprotein (P), ensures the transcription and the replication of the viral genome (Quarleri et al., 2022). Beyond the P protein, the P gene also drives the synthesis of the V, W, and C proteins. The V and W proteins are produced through a co-transcriptional editing mechanism involving the addition of one (V) or two (W) non-templated G nucleotides at the editing site of the P messenger (Escudero-Pérez et al., 2023). As a result, the P, V and W proteins share a common N-terminal domain (NTD), and have unique and distinct C-terminal domains (CTD). The W proteins of HeV and NiV contain a nuclear localization sequence (NLS) in their unique CTD leading to their nuclear accumulation in infected cells. Nuclear importation relies on binding to importins α3 and α4 and, to a lower extent, to importin α1 and α7 (Smith et al., 2018). W, along with P and V, also contains a nuclear export signal (NES) in its NTD ensuring its export to the cytoplasm of infected cells *via* a Crm-1-dependent nuclear export (Lo et al., 2009; Rodriguez et al., 2002). Since W is almost found in the nucleus of infected cells, the NLS overcomes the NES (Shaw, 2009; Tsimbalyuk et al., 2020).

The V and W proteins are virulence factors implicated in the inhibition of the antiviral response by binding several key cellular proteins (Basler, 2012; Gondelaud et al., 2022; Quarleri et al., 2022; Shaw, 2009; Wang et al., 2021). The lack of the V and W proteins in the non-pathogenic Cedar virus due to the absence of the RNA editing site, concomitantly with the absence of pathogenicity associated with this virus in humans, further supports the critical role of these proteins in pathogenicity (Laing et al., 2018; Marsh et al., 2012). W proteins bind and sequester STAT1 in the nucleus through their NTD inhibiting the interferon (IFN) response (Gondelaud et al., 2022; Quarleri et al., 2022; Shaw et al., 2004). Additionally, the CTD of the W proteins binds 14-3-3 proteins through its serine 449 leading to 14-3-3 protein nuclear accumulation. This accumulation enhances the nuclear export of p65, reducing its ability to interact with its promoter targets, thereby inhibiting the NF-κB-induced proinflammatory response (Enchery et al., 2021). Binding to 14-3-3 also provokes cell apoptosis through a less defined mechanism (Edwards et al., 2020). Presently, it is not known whether and how W proteins switch from inhibition of the IFN response to inhibition of the NF-κB-mediated response.

We previously showed that the W proteins from HeV and NiV (W^HeV^ and W^NiV^) are intrinsically disordered and can assemble to form highly flexible and curved fibrils (Pesce et al., 2022). W^NiV^ and W^HeV^ contain cysteine residues implicated in W dimerization (Pesce et al., 2022). We hypothesized that cysteine residues could impact the fibrillation ability of W proteins. Using various biophysical approaches and site-directed mutagenesis, we demonstrate here that the cysteine oxidation state dictates the ability of W^HeV^ and W^NiV^ to fibrillate and that nucleation relies on the formation of disulfide bridge-mediated dimers. We provide the first experimental evidence that W^HeV^ self-assembles also *in cellula* forming various types of condensates in the nuclei of transfected cells. These condensates differ in size and morphology and range from irregular, spherical and elongated assemblies to giant filamentous structures. In addition, they exhibit various dependencies on the redox state of cysteine residues. Our results also indicate that non-assembled W^HeV^ is more efficient at blocking STAT1-mediated IFN response while W^HeV^ redox-dependent condensates enhance the repression of the NF-κB-mediated antiviral response, suggesting a role in viral escape from the innate immune response.

## Results

### Cysteines are responsible for the formation of oligomeric species and of a compact conformation

The W^HeV^ and W^NiV^ proteins, which we previously showed to be able to form highly flexible and curved fibrils *in vitro* (Pesce et al., 2022), contain four and six cysteine residues respectively, three of which are shared by both proteins (**Figure 1A**). Since the W proteins are intrinsically disordered (Pesce et al., 2022) and thus possess a large solvent-accessible surface area, we hypothesized a strong reactivity of these residues. The proteins, purified in reducing and denaturing conditions, were incubated at 37°C in the presence of urea to avoid their spontaneous aggregation into fibrils while enabling cysteines to undergo natural oxidation, presumably *via* the formation of intra- and/or interchain disulfide bridges. This pre-incubation of the proteins in urea will thereafter be referred to as “oxidative preincubation” (**Supplementary Figure S1**).

**Figure 1.**
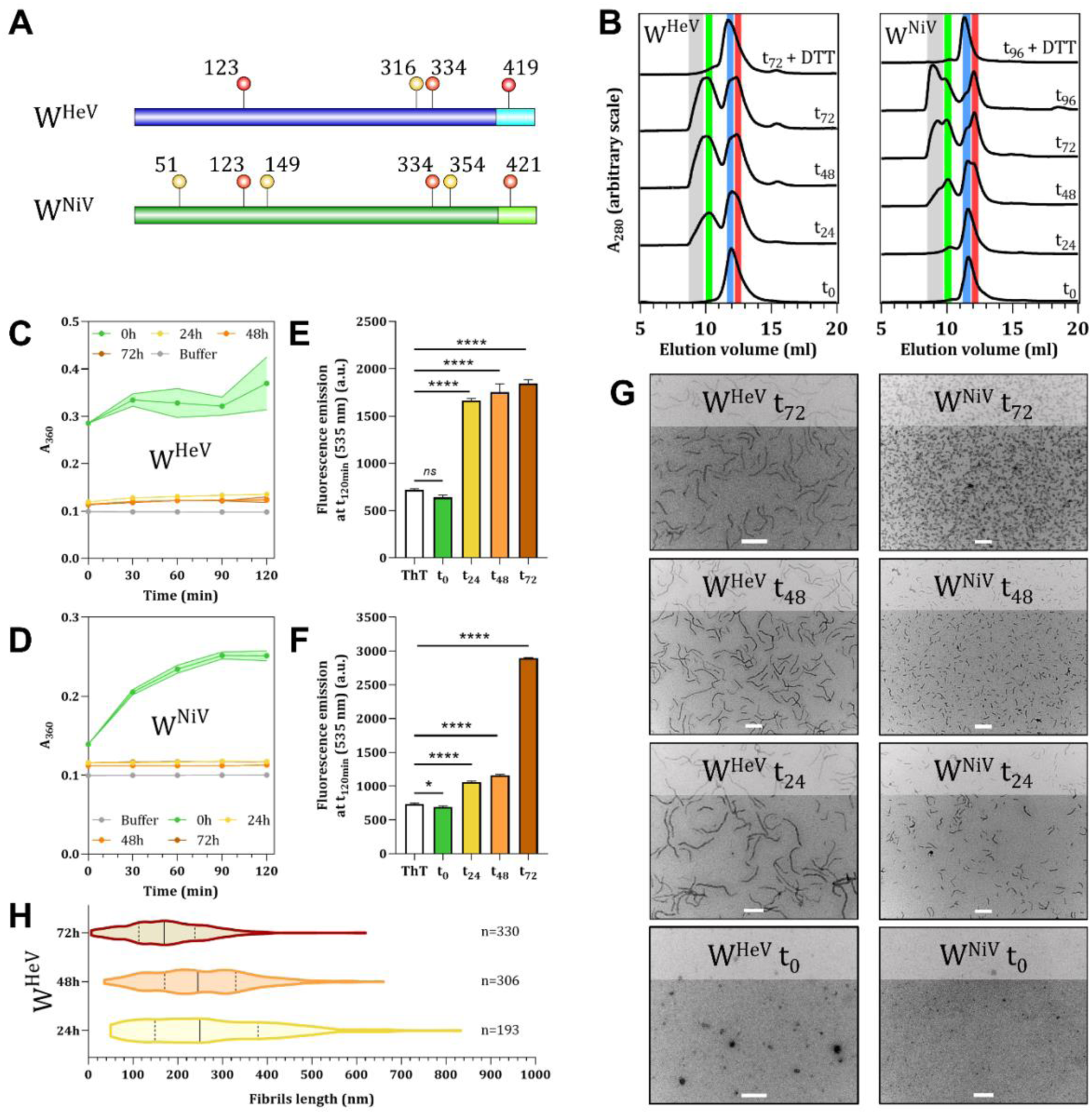
Cysteine residues oxidation of W^HeV^ and W^NiV^ is required to promote their fibrillation. (**A**) Location of cysteine residues in W^HeV^ (blue) and W^NiV^ (green) proteins. Conserved Cys are indicated in red. Dark colors: N-terminal domain (NTD), bright colors: C-terminal domain. (**B**) Analytical SEC of the W proteins after different preincubation times in oxidative condition at 37°C in the presence of urea, and either in the absence (t0, t24, t48, t72, and t96) or in the presence of DTT (t72 or t96 + DTT). The vertical grey, green, and blue bars correspond to the oligomeric, dimeric and monomeric species, respectively. The vertical red bar corresponds to a redox-dependent more compact monomeric form. (**C**) Turbidity and (**E**) ThT fluorescence measurements of W^HeV^ preincubated for 0 (green), 24 (yellow), 48 (orange), and 72h (brown) at 37°C in urea and then buffer exchanged to remove urea. Shown is also the fluorescence of ThT alone (white). Panels C and D show the average values and standard deviation (s.d.) as obtained from n=4 independent measurements. Statistical analysis was made with a one-way ANOVA, Dunnett’s test, ns: not significant, ****: p<0.0001. The shaded region in (C) corresponds to the error bar. (**D**) Turbidity and (**F**) ThT fluorescence measurements of W^NiV^ preincubated or not. The legend is the same as for (C) and (D). (**G**) NS-EM micrographs of W^HeV^ and W^NiV^ incubated for 120 min at 37°C in urea-free buffer without preincubation (0h) or following a preincubation of 24, 48 and 72h at 37°C in urea. (**H**) W^HeV^ fibril contour length measurements from NS-EM micrographs after 24, 48, and 72h of preincubation. Median and quartiles are represented as continuous and dotted lines, respectively. The number of fibril length measurements is indicated for each condition. Scale bar: 200 nm.

In addition to the monomer (vertical blue line in **Figure 1B** and peak I in **Supplementary Figure S2A**), size exclusion chromatography (SEC) analysis of the W proteins during oxidative preincubation showed the fast formation of dimeric species (vertical green line in **Figure 1B**), consistent with previous observations (Pesce et al., 2022), as well as the formation of higher-order oligomers within 24h for W^HeV^ and 48h for W^NiV^ (vertical grey line in **Figure 1B**). The nature of the oligomeric species of W^HeV^ was disentangled by mass photometry, which showed the co-existence in the sample of monomeric, dimeric, trimeric and likely tetrameric species (**Supplementary Figure S2B**). Additionally, a species (vertical red line in **Figure 1B** and peak II in **Supplementary Figure S2A**) that eluted at a volume slightly larger than that of the initial monomeric species can be detected upon incubation. For W^HeV^, non-reducing SDS-PAGE and mass spectrometry analysis revealed that this species corresponds to a monomeric form of the protein (**Supplementary Figures S2C** and **S2D**), and its elution volume indicates a more compact conformation (**Figure 1B**).

While W^HeV^ oligomerization does not seem to evolve further after 48h of oxidative preincubation, in the case of W^NiV^ it continues to progress even after 4 days (**Figure 1B**). Addition of DTT during the oxidative preincubation in urea (for 72h for W^HeV^ and for 96h for W^NiV^), led to the disappearance of both the oligomers (grey line in **Figure 1B**) and the compact conformation (red line in **Figure 1B**) indicating that they both are redox-sensitive and result from the formation of inter- and intramolecular disulfide bonds, respectively (**Figure 1B**).

### Cysteine oxidation is required for W^HeV^ and W^NiV^ fibrillation

The aggregation of W^HeV^ and W^NiV^ was followed by turbidity as well as by thioflavin T (ThT) fluorescence measurements (**Figure 1C-F**). In these experiments, as well as in all the ensuing experiments unless differently stated, the W proteins were pre-incubated for different periods in urea to promote the formation of oligomers as in **Figure 1B**. After this oxidative preincubation period, urea was removed and the aggregation process was monitored for 2h at 37°C in a urea-free buffer (see procedure in **Supplementary Figure S1**). Urea removal without an oxidative preincubation, *i.e.* at time 0, led to the formation of macroscopically visible insoluble aggregates (**Figure 1C** and **1D**) that eventually sedimented and that did not bind ThT (**Figure 1E** and **1F**). By contrast, oxidative preincubation of W proteins for at least 24h led to a clear solution (**Figure 1C** and **1D**) containing species that bind ThT (**Figure 1E** and **1F**), suggesting the formation of fibrillar structures. After one or two additional preincubation days (48h and 72h in **Figures 1C** and **1E**), both the turbidity and the fluorescence emission only poorly evolved for W^HeV^, coincidentally with the arrest of oligomerization as observed by analytical SEC (**Figure 1B**). In contrast, W^NiV^ ThT binding continued to gradually increase (**Figure 1F**), mirroring the evolution of its oligomerization (**Figure 1B**). The nature of the aggregates formed with or without an oxidative preincubation period was assessed by negative-staining electron microscopy (NS-EM). Grids were prepared after the 2h of incubation over which the aggregation and ThT-binding measurements were carried out (**Figure 1G**). The insoluble aggregates that are formed without an oxidative preincubation have an amorphous nature, while W^HeV^ and W^NiV^ preincubated in oxidative conditions give rise to fibrils (**Figure 1G**).

The constant increase of W^NiV^ oligomers content over time can be ascribed to its higher content in cysteine residues that likely keep reticulating the protein. The balance between fibril nucleation and elongation is low in the case of W^HeV^, as judged from the fact that it rapidly forms fewer, though longer fibrils than W^NiV^. By contrast, W^NiV^ exhibits a much higher nucleation, as judged from the significantly higher number of significantly shorter fibrils detectable on the micrographs taken at various times during the oxidative preincubation (**Figure 1G**). The ThT fluorescence intensity seems to correlate with the nucleation extent rather than with fibril size (*cf.* panels E and F with panel G of **Figure 1**).

W^NiV^ and W^HeV^ fibrils have the particularity of being not straight and rather highly curved (**Figure 1G**). Measurements of the contour length of W^HeV^ fibrils observed in micrographs revealed that 24h of oxidative preincubation led to the longest fibrils, up to 830 nm in these experimental conditions, with an average size of 273 nm (**Figure 1H**). The proportion of short fibrils increased with the incubation time, presumably reflecting fibril fragmentation that regularly occurs in amyloid fibrils. After 72h of incubation, the average length of fibrils dropped to 179 nm (**Figure 1H** and **Supplementary Table 1**).

Taken together, these observations suggest that disulfide bonds formation is the initial step in the fibrillation of both W^HeV^ and W^NiV^ proteins, since non-oxidized monomeric species are poorly able to undergo fibrillation. To ascertain the hypothesis that oligomers promote the formation of fibrillar structures, we focused on W^HeV^ in light of its lower cysteine content and its faster oligomerization compared to W^NiV^. Since 48h of oxidative preincubation of W^HeV^ represents a good compromise between solubility and fibril length, and because the oligomerization process does not progress further after this period, this preincubation time was selected for the following analyses of W^HeV^ fibrillation.

The involvement of cysteine residues in W^HeV^ fibrillation was confirmed by generating a variant where the four cysteine residues were all substituted to serine residues (W^HeV^ C^all^S) (**Figures 2A-D**). As expected, W^HeV^ C^all^S remains monomeric after 48h of oxidative preincubation in urea (**Figure 2A**). After urea removal, insoluble aggregates that do not bind ThT were detected as in the case of the wild-type (WT) protein not subjected to the oxidative preincubation (**Figure 2B** and **2C**). Under these conditions, NS-EM confirmed the sole presence of amorphous aggregates (**Figure 2D**). Furthermore, the addition of DTT on preincubated W^HeV^ immediately after urea removal, *i.e.,* at t_0_, led to a progressive decrease in solubility over the 120 min of incubation (**Figure 2B**). After 120 min, ThT binding was weaker in the presence of DTT (red bar) than in its absence (orange bar), but not as low as that observed with the non-incubated WT protein (green bar) or with the W^HeV^ C^all^S variant (blue bar) (**Figure 2C**). NS-EM micrographs taken after 120 min of incubation in the presence of DTT showed the co-existence of rare and small fibrils, and of amorphous aggregates (**Figure 2E**).

**Figure 2.**
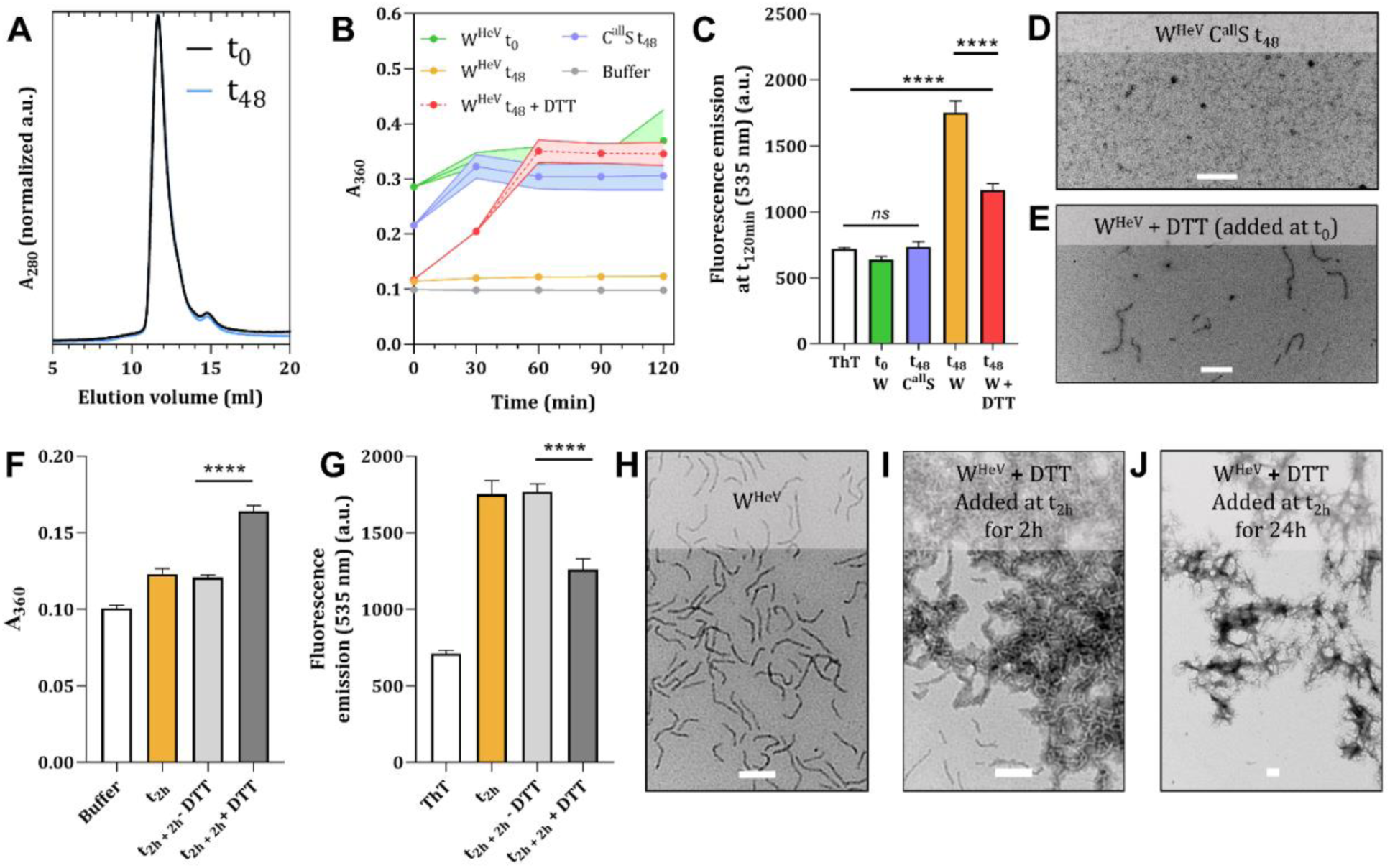
Cys-to-Ser substitution abrogates W^HeV^ fibrillation and aggregation in fibrillar structures cannot be reversed by reduction. (**A**) Analytical gel filtration of W^HeV^ C^all^S preincubated for 48h (t48h, blue) or not (t0h, black) in urea. (**B**) Turbidity and (**C**) ThT fluorescence measurements of W^HeV^ C^all^S (blue) and of the WT W^HeV^ protein in the presence of DTT (red), both preincubated for 48h. Non-incubated (0h, green) and preincubated for 48h (orange) WT W^HeV^ data are shown for comparison. NS-EM micrographs were taken after 2h of incubation of W^HeV^ C^all^S preincubated for 48h in urea (**D**) and of WT W^HeV^ in the presence of DTT added after urea removal (**E**). (**F**) Turbidity and (**G)** ThT fluorescence measurements of W^HeV^ preincubated for 48h and after 2h of incubation following urea removal (orange), and after 2 extra hours in the presence (dark grey) or absence (light grey) of DTT. NS-EM micrographs of W^HeV^ preincubated for 48h in urea and after 2h of incubation following urea removal *plus* 2 extra hours in the absence (**H**) or presence (**I**) of DTT. W^HeV^ was preincubated for 48h in urea and then for 24h in urea-free buffer in the presence of DTT. The solution was centrifuged and the resuspended pellet (**J**) was observed by NS-EM. Panels C and G also display the fluorescence of ThT alone (white). Panels B, C, F, and G show the average values and error as obtained from n=4 independent measurements. Statistical analysis was made with a Student’s t-test, ns: not significant, ****: p<0.0001. The shaded region in B) corresponds to the error bar. Scale bar: 200 nm.

To further confirm the relation between oligomerization and fibrillation, we artificially generated oligomers by cross-linking W^HeV^ C^all^S using DTSSP (**Supplementary Figure S3**), a crosslinker containing a disulfide bond that thus drives the formation of disulfide-bridged oligomers mimicking those formed by the WT protein. Upon crosslinking, W^HeV^ C^all^S was found to be remarkably soluble during the aggregation measurements (**Supplementary Figure S3A**), to strongly bind ThT (**Supplementary Figure S3B**) and to fibrillate after 2h of incubation (**Supplementary Figure S3C**).

### Formation of W^HeV^ fibrillar structures is not reversible upon reduction

To assess whether the fibrillation process was reversible, W^HeV^ was incubated 48h in oxidative conditions thus enabling disulfide bond formation, and then allowed to assemble into amyloid-like fibrils for 2h after urea removal, before the addition of DTT (**Figure 2F-J**). Aggregation measurements and NS-EM were performed after two extra hours (t_2h + 2h_) following DTT addition. As expected, these experiments showed decreased solubility and ThT binding (**Figure 2F** and **2G**) and the presence of massive clusters of fibrillar structures that were not observed in the absence of DTT (**Figure 2**, compare **H** and **I**).

After 24h of incubation in the presence of DTT, the samples were centrifuged and the pellet was resuspended and deposited on an NS-EM grid (**Figure 2J**). NS-EM showed large filamentous structures of micrometric scale consisting of clustered fibrils (**Figure 2J**) indicating that the fibrillation process is not fully reversible through reduction, though a limited fraction of W^HeV^, presumably monomers detached from the fibrils, became insoluble and amorphous according to the aggregation measurements made just after 2h of incubation. Thus, while disulfide-bridged oligomers and/or the redox-sensitive compact conformation, *i.e.,* species II in **Supplementary Figure S2A**, play a critical role in fibrils nucleation, once fibrils have formed, a redox-independent mechanism maintains monomers together. Disulfide bonds also regulate fibril assembly while their reduction in mature fibrils causes random association of W^HeV^ leading to the formation of fibrillar clusters.

### Dimeric species are sufficient to initiate fibrillation

In order to identify the minimal molecular assembly required for fibril nucleation, W^HeV^ triple Cys to Ser variants were generated, and SEC analysis showed that they are only able to dimerize, although in different proportions (**Figure 3A**). After 48h of preincubation in oxidative conditions and a 2h-incubation after urea removal, the four variants were shown to be able to bind ThT (**Figure 3B**) and to fibrillate (**Figure 3C**) indicating that the sole presence of dimeric species is sufficient to promote W^HeV^ fibrillation. At first glance, the nucleation extent, *i.e.* the number of fibrils observed on micrographs, seems to be at least partly related to the initial amounts of dimers observed in SEC analysis. Indeed, the W^HeV^ C123^only^, C316^only^, and C419^only^ variants, which display considerable amounts of dimers, give rise to micrographs characterized by a high density of relatively short fibrils. Conversely, the W^HeV^ C334^only^ variant, which weakly dimerizes, displays attenuated nucleation as judged from the presence of rare fibrils in micrographs (**Figure 3C**). These observations strongly support the implication of dimeric species in the nucleation step of W^HeV^ fibrillation.

**Figure 3.**
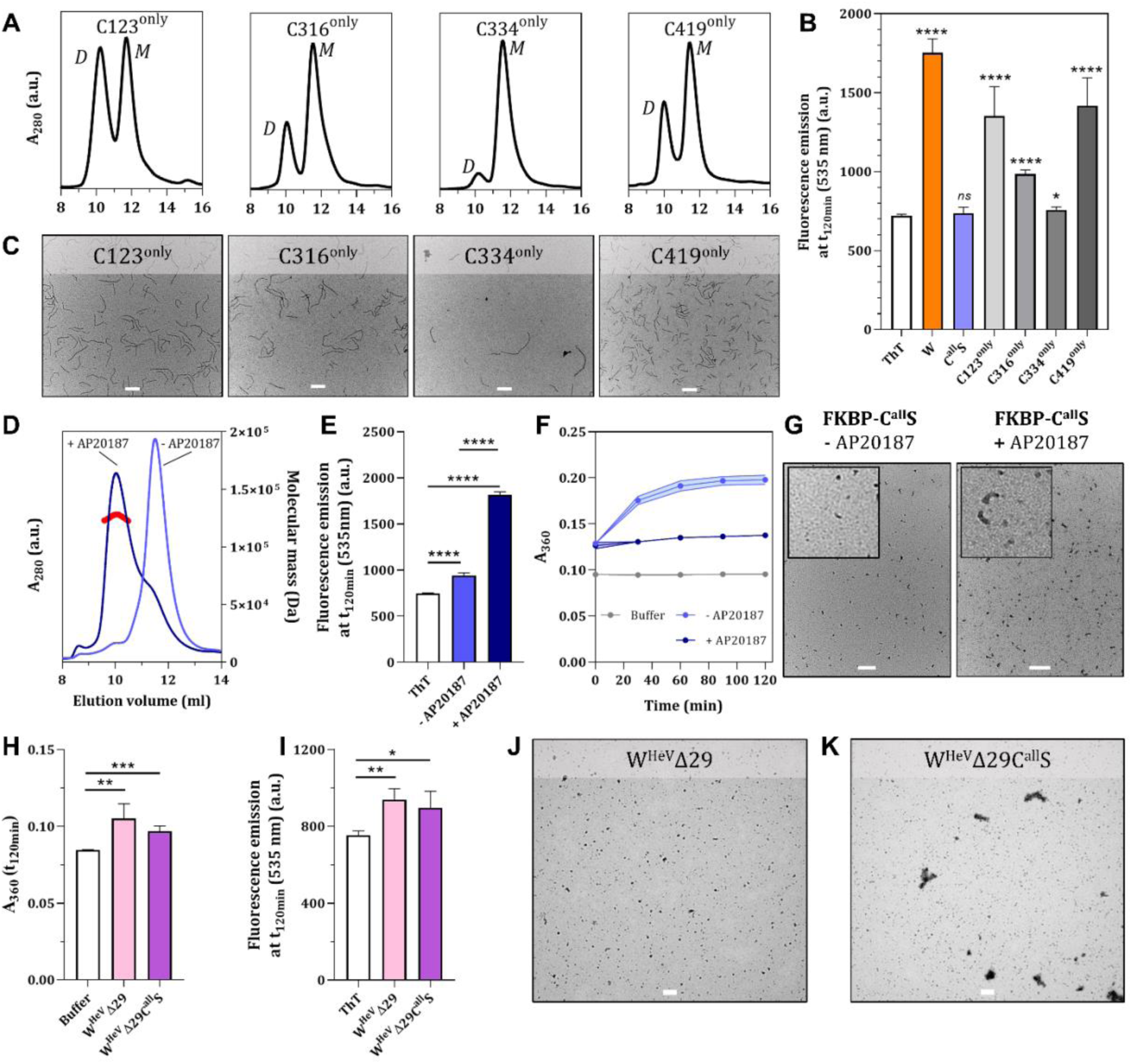
W^HeV^ dimerization is sufficient to promote fibrillation and the 2-29 region is strictly required for fibrillation. (**A**) Analytical SEC of W^HeV^ with triple Cys to Ser substitutions preincubated 48h in urea. M and D stand for monomeric and dimeric species, respectively. (**B**) ThT fluorescence measurements of W^HeV^ triple variants, preincubated in urea for 48h, after 2h in urea-free buffer. Shown are mean values and s.d. as obtained from n ≥ 4 independent measurements. Panel B also shows the fluorescence of ThT alone (white) and of W^HeV^ (orange) and W^HeV^ C^all^S (blue) for comparison. Statistical analyses were performed using the Student’s t-test, comparing fluorescence data to those obtained with ThT alone. (**C**) NS-EM micrographs of W^HeV^ with triple Cys to Ser after 2h of incubation following urea removal. Scale bar: 200 nm. (**D**) Analytical SEC of FKBP-C^all^S in the presence (dark blue) or absence (light blue) of the AP20187 molecule along with the molecular mass determination by SEC-MALLS of FKBP-W^HeV^ C^all^S in the presence of AP20187 (127 800 Da, red). Turbidimetry (**E**) and ThT fluorescence measurements (**F**) of FKBP-W^HeV^ C^all^S preincubated (dark blue) or not (light blue) with AP20187. Panel E also shows the fluorescence of ThT alone (white). (**G**) Micrographs of FKBP-C^all^S after 2h of incubation with or without AP20187. Inset: zoom. Panels B, E, and F show mean values and s.d. as obtained from n=4 independent measurements. The shaded region in F) corresponds to the error bar. Statistical analysis was made with a Student’s t-test; ns: not significant, ****: p<0.0001. Scale bar: 200 nm. (**H**) Turbidity and (**I**) ThT fluorescence measurements of W^HeV^Δ29 (pink) and W^HeV^Δ29 C^all^S (violet) preincubated for 48h at 37°C in urea and then for 2h in urea-free buffer. Shown is also the fluorescence of ThT alone (white). Panels H and I show the average values and standard deviation (s.d.) as obtained from n=4 independent measurements. Statistical analysis was made with Student’s t-test, *: p<0.05, **: p<0.01, ***: p<0.001. NS-EM micrographs of W^HeV^Δ29 (**J**) and W^HeV^Δ29 C^all^S (**K**) incubated for 120 min at 37°C in urea-free buffer following a preincubation of 48h at 37°C in urea.

Finally, we produced dimeric species by generating a protein construct (FKBP-C^all^S) whose dimerization can be chemically induced (**Figure 3D-G**). FKBP-C^all^S was made by fusing W^HeV^ C^all^S and FK506-binding protein, FKBP. FKBP is a widely-used system to study homodimerization processes since the protein strongly dimerizes in the presence of the AP20187 molecule (DeRose et al., 2013). FKBP-C^all^S is expected to be unable to fibrillate in the absence of AP20187, while the addition of the latter should promote the nucleation of fibrils *via* the formation of a large amount of chemically-induced dimeric species. After 10 minutes of incubation in the presence of AP20187, analysis of FKBP-C^all^S by SEC-MALLS showed that the chimeric protein was effectively predominantly dimeric (**Figure 3D**). Urea and unbound AP20187 were removed and the aggregation and binding to ThT were monitored for 2h (**Figure 3E** and **3F**). In these experiments, and in line with expectations, FKBP-C^all^S turned out to be more soluble and to bind ThT to a higher extent in the presence of AP20187 than in its absence (**Figure 3E** and **3F**). In the absence of AP20187, the protein slightly binds ThT, an observation that can be accounted for by the initial presence of low amounts of dimeric species in the purified sample. Consequently, a very small number of very short fibrils was observed without AP20187, while their number greatly increased in its presence (**Figure 3G**). Because of the large molar excess of AP20187 used in these experiments, dimerization is strongly promoted thereby favoring fibril nucleation at the expense of elongation, resulting in numerous and very short fibrils (**Figure 3G**).

Taken together, these observations demonstrate that W^HeV^ dimerization is the driver of the nucleation process and represents a critical step in the formation of fibrils. A similar conclusion can be drawn for W^NiV^ owing to the correlation between the W^NiV^ oligomers proportion and its nucleation extent (see **Figure 1**).

### The N-terminal region of W^HeV^ spanning residues 2 to 29 is strictly required for fibrillation

We recently reported that the N-terminal region encompassing residues 2 to 110 of the W^HeV^ protein (referred to as PNT1) is the most fibrillogenic region among three NTD fragments (PNT1 to PNT3) collectively covering residues 2 to 310 (Gondelaud et al., 2025). A predicted cryptic amyloidogenic region (CAR, aa 10 to 19) within PNT1 was found to be the main driver of fibrillation, with the downstream region encompassing residues 20 to 29 contributing to the process (Gondelaud et al., 2025). Removal of the 2-29 region in the context of PNT1 fully abrogated fibrillation (Gondelaud et al., 2025). Here, we generated a W^HeV^ variant devoid of the first 29 residues (referred to as W^HeV^ Δ29) as well as the W^HeV^ Δ29C^all^S variant that also lacks cysteine residues. Following an oxidative preincubation, that is, a condition enabling W^HeV^ to form fibrils, and after an additional 2h incubation in the absence of urea, both variants were found to be slightly insoluble (**Figure 3H**), to poorly bind ThT (**Figure 3I**) and to be unable to fibrillate (**Figure 3J, K**), with micrographs showing only a large number of small assemblies.

Collectively, these data indicate that although oxidized cysteines are necessary, they are not sufficient for W^HeV^ fibrillation to take place. Results enable mapping to residues 2-29 the region strictly required for W^HeV^ fibrillation.

### W^HeV^ forms different types of condensates in the nuclei of transfected cells

We next assessed the ability of W^HeV^ to also form fibrils *in cellula*. To allow the visualization of W^HeV^ in living cells (*i.e.*, non-fixed and non-permeabilized cells), we generated constructs driving the eukaryotic expression of W^HeV^ fused to a tetracysteine (*tc*) peptide tag (Arhel and Charneau, 2009; Martin et al., 2005) appended either at the N- (*tc*-W) or C-terminus of the protein (W-*tc*). Human embryonic kidney cells (HEK293T) were transfected with these constructs or with an empty control plasmid (EV) or with a plasmid driving the expression of the enhanced green fluorescent protein (EGFP). W^HeV^ was detected by confocal microscopy at 24h post-transfection using either the membrane-permeable ReAsH dye, which fluoresces upon binding to the *tc* tag, or by immunofluorescence (IF) staining targeting W^HeV^.

In some transfected cells expressing W^HeV^, nuclear filamentous structures along with irregular aggregates could be observed both at 24h and 48h post-transfection (**Figures 4A**, **B**). Notably, *tc* staining (red) was only visible in W^HeV^-stained nuclei (magenta), confirming that the filaments are composed of W^HeV^ (**Figure 4A**). These structures were detectable neither in cells transfected with the empty vector, nor in cells expressing EGFP (**Figure 4A**), thus ruling out the possibility of non-specific ReAsH binding. The filamentous structures and irregular aggregates could also be detected upon staining with FlAsH, another *tc*-specific dye displaying higher contrast and less cytotoxicity in our hands (**Figure 4C**). Of note, filamentous structures and irregular aggregates were observed in both HEK293T and BEAS-2B cells, the latter being human bronchial epithelial cells that constitute a relevant cellular model for henipavirus infection that always targets the respiratory tract (**Figure 4C**).

**Figure 4.**
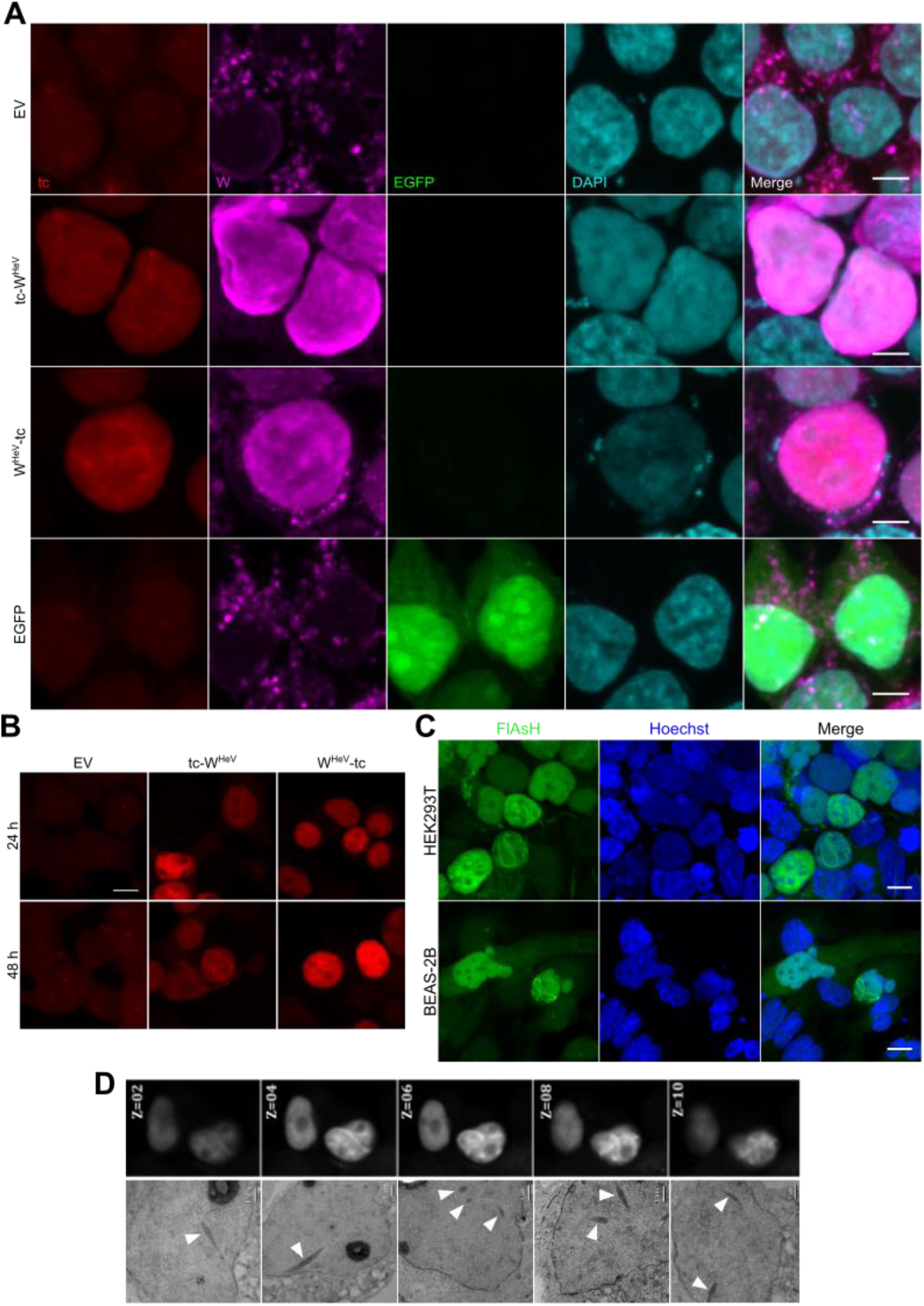
W^HeV^ forms filamentous nuclear structures in cellula. (**A**) Immunofluorescence and confocal microscopy analysis of HEK293T cells transfected with either an empty vector (EV), or enhanced green fluorescent protein (EGFP), or the indicated tetracysteine (tc)-tagged W^HeV^ constructs. W^HeV^ filaments were visualized 24h post-transfection with ReAsH dye to detect the tc peptide (red) and immunofluorescence staining targeting W^HeV^ (magenta). Nuclei were counterstained with DAPI (blue). Scale bar, 10 µm. (**B**) Z-stack projections of live HEK293T cells transfected with an empty vector or the indicated tc-tagged W^HeV^ constructs. Cells were stained 24h and 48h post-transfection with ReAsH dye to detect the *tc* peptide and imaged by confocal microscopy. Images are representative of three independent experiments. Scale bar, 5 µm. (**C**) Z-stack projections of HEK293T and BEAS-2B cells transfected with the *tc*-W^HeV^ construct. Cells were stained 48h post-transfection with FlasH dye and Hoechst. Scale bar, 10 µm. (**D**) Z-stack projections of HEK293T cells transfected with the W^HeV^-tc construct, stained and imaged at 24h post-transfection as described in panel B (top). Corresponding electron micrographs of ultrathin sections at equivalent Z positions of the cells shown in top confocal images. The arrowheads indicate the fibrillar structures detected in the nuclei of the transfected cells (bottom).

EM analysis of ultrathin sections of transfected HEK293T cells expressing W-*tc* at 24h post-transfection and displaying ReAsH-stained nuclear filaments, confirmed the presence of nuclear filamentous structures (**Figure 4D)**. These structures were consistently observed at several *z* stacks of the reference fluorescence stack (**Figure 4D**, see Material and Methods and SI for details of the correlative approach). Similar structures were also observed at 48h post-transfection (**Supplementary Figure S4A**), while they were never observed in non-transfected cells (**Supplementary Figure S4B)**, nor in transfected cells in which no filamentous structures could be detected by confocal microscopy (**Supplementary Figure S4C**).

Noteworthy, the nuclear filaments appear to be at least 10 times longer than the fibrils observed *in vitro*, being in the micrometer, rather than in the sub-micrometer, length scale (cf. **Figure 1H** and **Figure 4**). Unexpectedly, *tc*-W is significantly more prone to form filaments compared to W-*tc* indicating that the *tc* tag interferes with the ability of W^HeV^ to form filaments when present at the C-terminus of the protein (**Supplementary Table 2**). The expression level of W^HeV^ variants was verified by IF staining and appeared to be higher for *tc*-W^HeV^ compared to W^HeV^-*tc* constructs (**Supplementary Figure S5**), a finding that can be regarded as another plausible explanation for the reduced propensity to form filaments of W^HeV^-*tc* constructs. In light of this, we focused on the *tc*-W^HeV^ construct for subsequent analysis, and switched from the pcDNA3.1(+) plasmid backbone to a pCAGGS backbone to drive higher expression levels. Unless differently stated, experiments described in the following sections were carried out on HEK293T cells.

Beyond filaments with different morphologies, W^HeV^ also forms other types of nuclear condensates (**Figure 5**). Specifically, at 24h and 48h post-transfection, W^HeV^ forms condensates of globular and more elongated shape (hereafter referred to as *spots* and *rods*, respectively) as well as more irregular aggregates (hereafter referred to as *aggregates*) (**Figure 5A, B**). The various types of condensates and the filaments are not mutually exclusive. Indeed, they are detectable concomitantly in some cells, as illustrated by the co-existence of filaments with irregular aggregates (see black arrows in **Figure 5A**), or spots with rods (see white asterisk in **Figure 5A**). A similar pattern was observed in BEAS-2B cells (**Figure 5C**) and in human lung carcinoma A549 cells (**Supplementary Figure S6**). Note that no condensates were observed in cells transfected with a control *tc*-mStayGold construct (**Figure 5D**).

**Figure 5.**
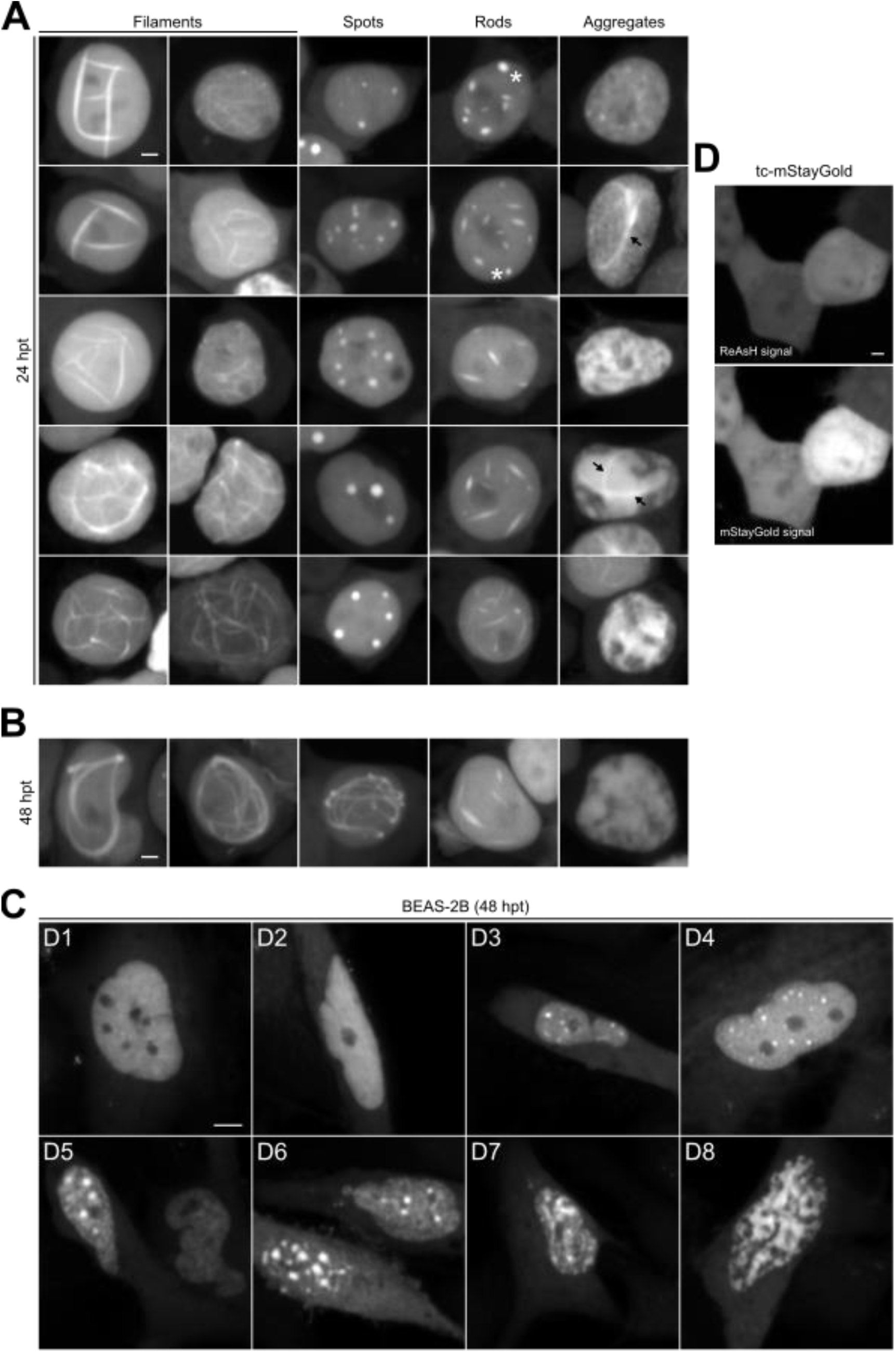
W^HeV^ forms different types of condensates in the nucleus of transfected cells. (**A, B**) Confocal microscopy imaging of HEK293T cells transfected with *tc*-W^HeV^ construct and stained with ReAsH at 24h (**A**) and 48h post-transfection (**B**). Filaments, spots (globular condensates), rods (short straight structures), and irregular aggregates (sometimes containing filaments, shown by arrows) are detectable in the nuclei. Scale bars, 2.5 µm. (**C**) Confocal microscopy imaging of BEAS-2B cells transfected with *tc*-W^HeV^ and stained with ReAsH at 48h post-transfection. Homogenous, diffuse nuclear W (D1, D2), rods (D3), spots (D3, D4, D5, D6), and amorphous aggregates (D7, D8) are detectable in the nuclei. Scale bar, 5 µm. (**D**) Confocal microscopy imaging of HEK293T cells transfected with a control *tc*-mStayGold construct and stained with ReAsH at 24h post-transfection, showing the absence of condensates, and only homogenous, diffuse signal. Scale bar, 2.5 µm.

In an attempt at unveiling the kinetics of condensate formation, *i.e*., the temporal relationships among the different types of condensates, we performed time-lapse confocal microscopy analyses of transfected HEK293T cells and used the milder FlAsH reagent to preserve cellular integrity (**Figure 6** and **Supplementary Figure S7**). Results support a stochastic scenario for condensate formation: filaments can form from an initially homogeneous and diffuse W signal (**Figure 6A**), and can then turn into irregular aggregates or seemingly disappear (**Figure 6B** and **Supplementary Figure S7A**), and spots can form from an initially homogeneous W signal (**Figure 6C**). In other words, and rather unexpectedly, rods and filaments do not correspond to different stages of a “*maturation*” process from spots, and irregular aggregates do not appear to be the last stage of the condensation process. Interestingly, some nuclei display a homogeneous W signal that intermittently transitions into irregular aggregates before reverting to a uniform distribution (**Supplementary Figure S7B**, white arrows), indicating that the amorphous aggregation of W is reversible. Overall, our data also highlight that the condensates and especially the filaments are highly dynamic, in that they adopt various conformations throughout the imaging time (**Figure 6** and **Supplementary Figure S7A** and **S7C)**.

**Figure 6.**
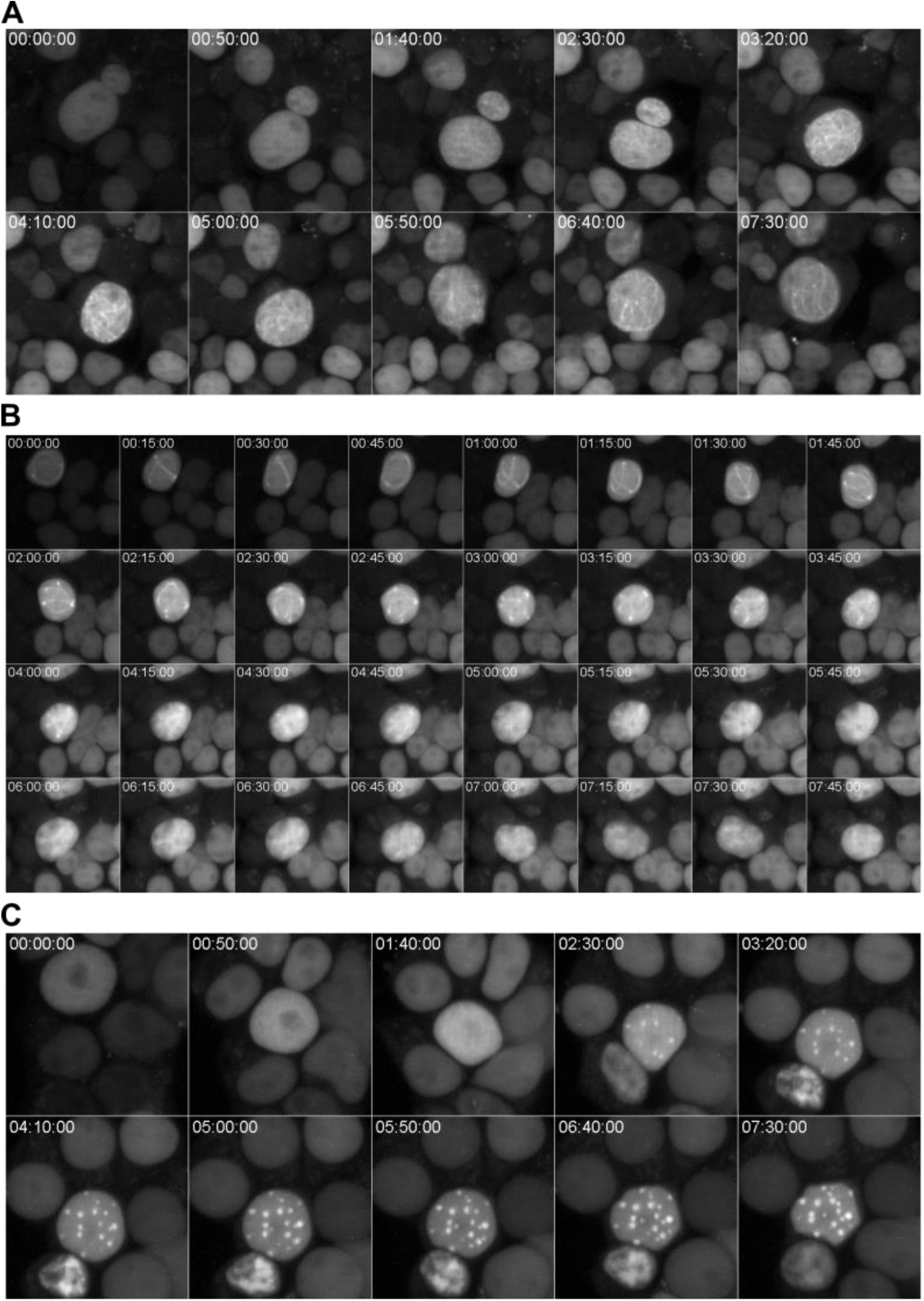
Time-lapse imaging of transfected HEK293T cells showing the formation and dynamics of W filaments and of the other types of condensates. (**A-C**) Z-stack projections of live HEK293T cells transfected with *tc*-W^HeV^ construct, and imaged by confocal microscopy in the presence of FlAsH, at the indicated times (in the format h:min:sec). (**A**) Appearance of filamentous structures in the nucleus from an initially homogeneous W signal. (**B**) Evolution of the morphology of a nuclear filament, which seems to turn into an amorphous structure. (**C**) Appearance of spots in the nucleus from an initially homogeneous W signal.

Based on high-resolution confocal microscopy acquisitions obtained with transfected HEK293T cells over different Z optical sections (**Figure 7A** and **Supplementary Movies 1 & 2**), the tridimensional (3D) reconstruction of both the globular condensates and the filaments was reconstituted and revealed various morphologies (**Figure 7B**). These high-resolution acquisitions and the subsequent condensates segmentation using Imaris software distinctly showed globular spots with the presence of amorphous aggregates (**Figure 7A1, B**), and long and intertwined filaments accompanied or not by irregular aggregates (**Figure 7A2-A5, B**). For filament-bearing nuclei, 3D reconstruction and segmentation indicate that only one filament with ramifications is present per nucleus (**Figure 7B**). Quantitative analysis indicated that the globular condensates and the filaments occupy as much as ∼5% of the volume of the nucleus, and that filaments have a section diameter of about 1 to 2 µm (**Figure 7B, C**).

**Figure 7.**
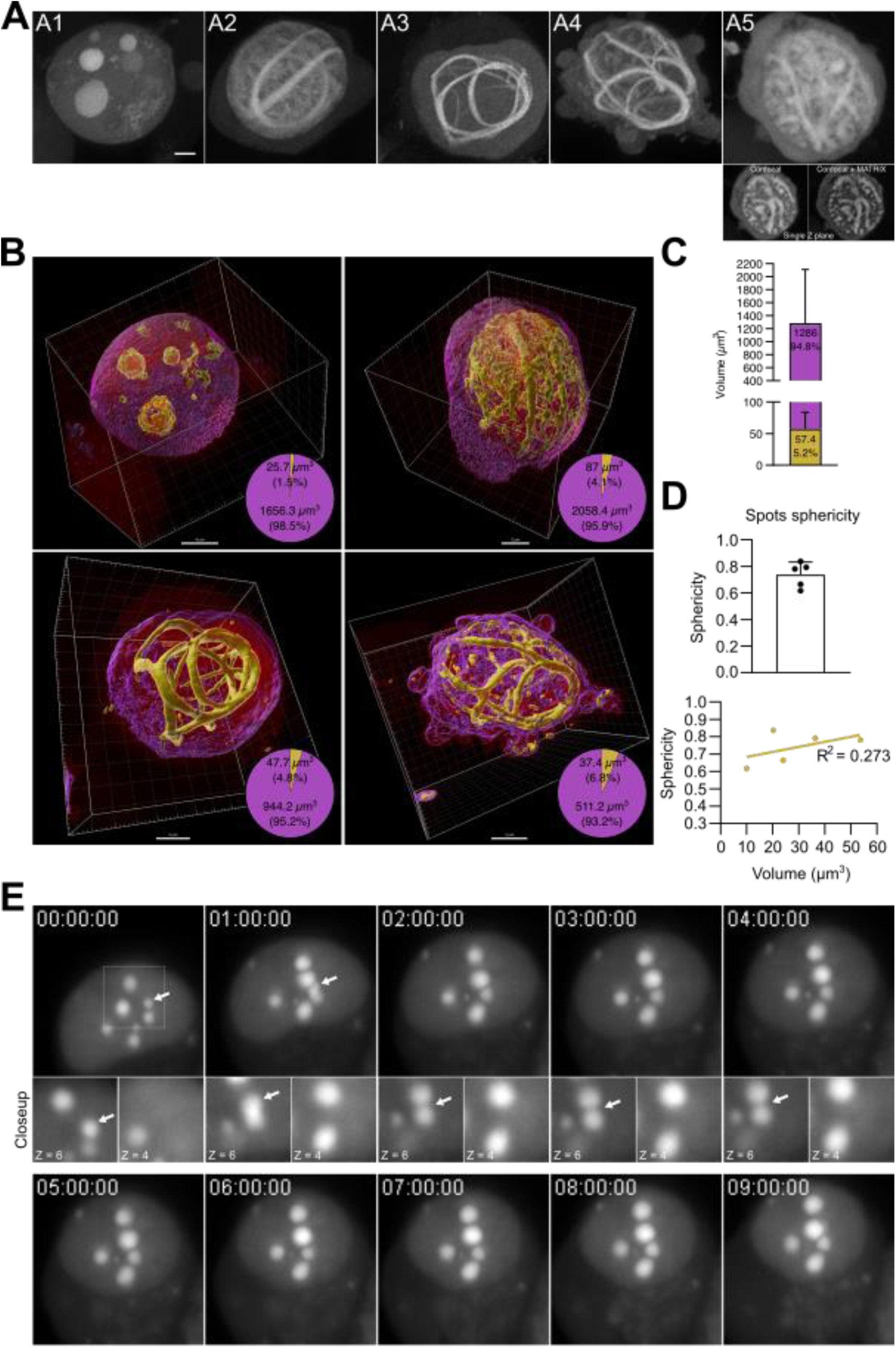
Tridimensional microscopy analysis of W filaments and condensates in transfected HEK293T cells. (**A**) Z-stack projections of live HEK293T cells transfected with *tc*-W^HeV^ construct, stained with ReAsH and imaged by high-resolution confocal microscopy at 72h post-transfection. Scale bar, 2.5 µm. (**B, C**) Segmentation, visualization and volume quantification of W structures with Imaris. The pie charts show the volume of the condensates (yellow) and the volume of the nuclei minus the volume of the condensates (violet). (**C**) Histogram indicating the mean volume of the filaments (yellow) and nuclei shown in panel B. (**D**) Analysis of the sphericity of the spots shown in panel B. (**E**) Time-lapse imaging of transfected HEK293T cells showing the dynamics and partial merging followed by segregation (arrows and closeup of single Z planes) of W spots. Indicated times are in the format h:min:sec.

We analyzed the sphericity of the spots to ascertain their possible liquid-like nature (**Figure 7D**). In the field of liquid–liquid phase separation (LLPS), sphericity is commonly used as a quantitative measure to assess whether a biomolecular condensate behaves like a liquid droplet. A sphericity value close to 1 is indicative of a liquid-like condensate, as liquids tend to minimize surface area due to surface tension, forming near-perfect spheres, while values <0.9 suggest solid-like, gel-like, or irregular morphology (Hyman et al., 2014). Obtained values support a non-liquid behavior of the spots (**Figure 7D**), and the lack of correlation between sphericity values and spot volume rules out the possibility that spots become more solid-like as their size increases (**Figure 7D**). In further support of a non-liquid nature, time-lapse imaging of the spots showed that they exhibit poor dynamics, *i.e*. they tend to occupy the same position over time, and also poorly coalesce (**Figure 7E**) (with coalescence being another hallmark of liquid-like behavior). Indeed, we did not detect any complete coalescence event, but only a few examples of partial merging of spots that eventually segregate again, a behavior consistent with a solid-like nature (**Figure 7E, arrows and closeup**).

### The various condensates formed by W^HeV^ are not associated with cell death

We next investigated the possible impact of the various types of condensates formed by W^HeV^ on cell death and viability (**Figure 8**). To this end, we stained cells with either propidium iodide or activated Caspase-1. Propidium iodide is a fluorescent dye that stains DNA in cells with compromised plasma membranes, and is therefore commonly used as a marker for dead or dying cells. Activated caspase-1 is primarily used as a marker of pyroptosis, a form of programmed cell death associated with inflammation. None of the cells exhibiting W^HeV^ condensates was found to be stained by propidium iodide or caspase-1 (**Figure 8A**) indicating that the various condensates induce neither cell death nor pyroptosis. Cells transfected with an irrelevant *tc*-mStayGold construct or treated with Triton X-100 served as negative and positive control, respectively. By contrast, alamarBlue metabolic assay, used as a proxy for cellular viability, revealed that transfection with the W^HeV^ expression plasmid was associated with a ∼30% reduction in metabolic activity compared to cells transfected with an empty vector (**Figure 8B**). Collectively, these data suggest that the observed decrease in cell viability does not reflect cell death, rather a general metabolic perturbation or reduced cell proliferation following ectopic expression of W.

**Figure 8.**
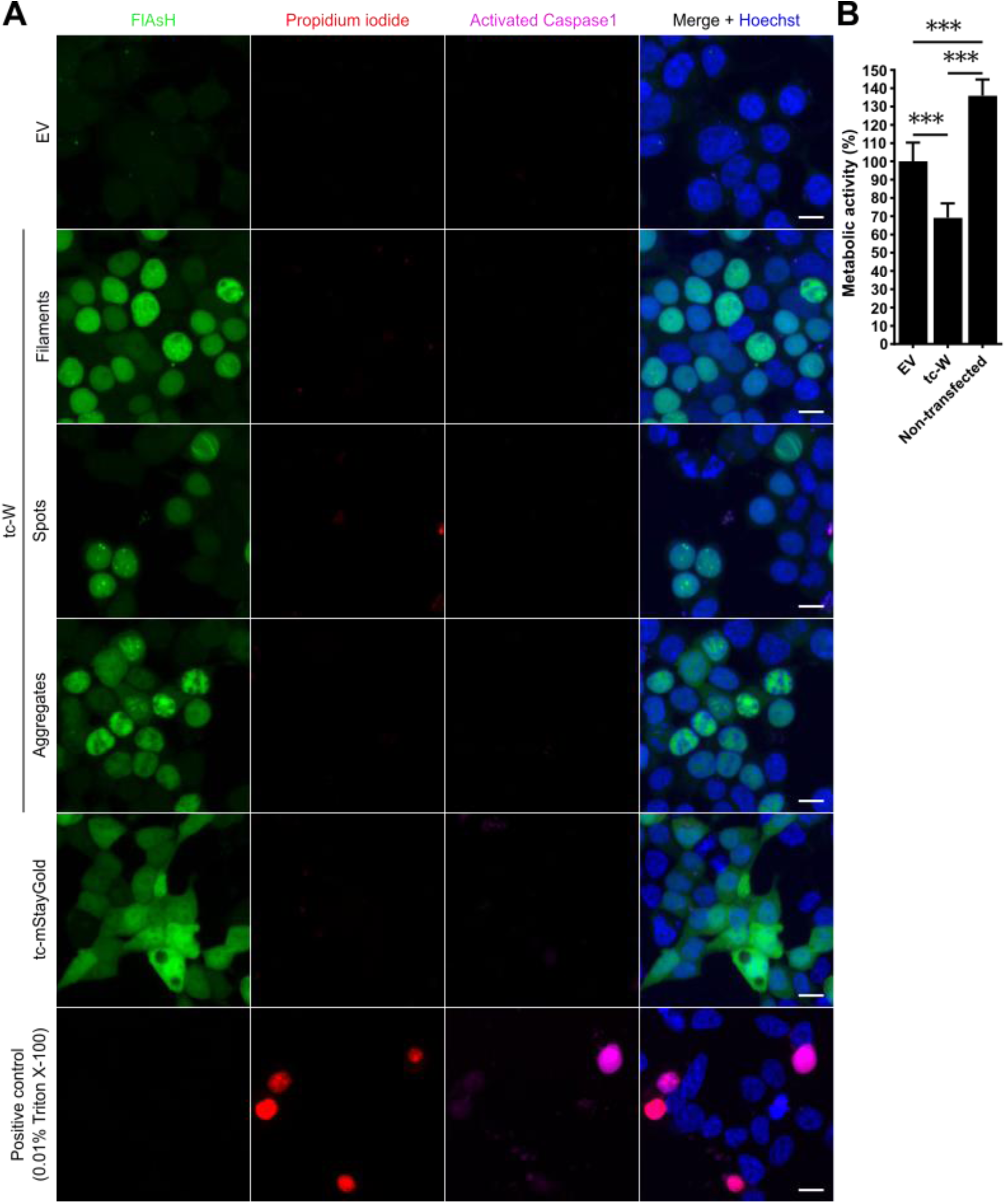
Neither filaments nor the other types of condensates formed by W^HeV^ are associated with cell death features. **A)** Z-stack projections of live HEK293T cells transfected with the indicated constructs, or only treated with 0.01% Triton X-100, and stained with FlAsH, propidium iodide, Hoechst and for activated Caspase-1, and imaged by spinning disk confocal microscopy at 24h post-transfection. Scale bar, 10 µm. **B)** Analysis of metabolic activity of HEK293T transfected cells with the indicated constructs by fluorometric alamarBlue assay, at 24h post-transfection. Data are expressed as mean ± sd and analyzed by one-way ANOVA completed by a Tukey’s multiple comparison test \*\*\**p*<0.001.

### Removal of the first 29 residues of W^HeV^ abrogates formation of filaments *in cellula*, while cysteine to serine substitution impairs the formation of the other types of condensates

To assess to which extent the phenotype of W^HeV^ variants observed *in vitro* translates into the cellular context, HEK293T cells were transfected with W^HeV^ constructs driving the expression of N-terminally *tc*-tagged W^HeV^ variants in which either all cysteines are replaced with serines (W^CallS^ variant), or the first 29 residues are deleted (WΔ29 variant), or both the deletion and the substitutions are combined (WΔ29^CallS^ variant) (**Figure 9A, B**). A construct encoding the WT form of the W^HeV^ protein was used for comparison. Quantitative analysis of the condensates formed by the different variants showed that WΔ29 does not form any filament at 24h post-transfection (**Figure 9C**). This observation mirrors the inability of this variant to form fibrils *in vitro*, thus suggesting a potential relationship and/or continuity between fibrils and filaments in spite of the large difference in size. We cannot exclude the possibility that the filaments result from the growth of fibrils and that the latter may escape detection *in cellula* due to confocal microscopy limitations in resolution. At 48h post-transfection, some cells, although at a very low proportion, display nuclear filaments, possibly suggesting that the cellular context might have a role in filamentation, providing potential additional drivers and pathways for filamentation compared to the *in vitro* context.

**Figure 9.**
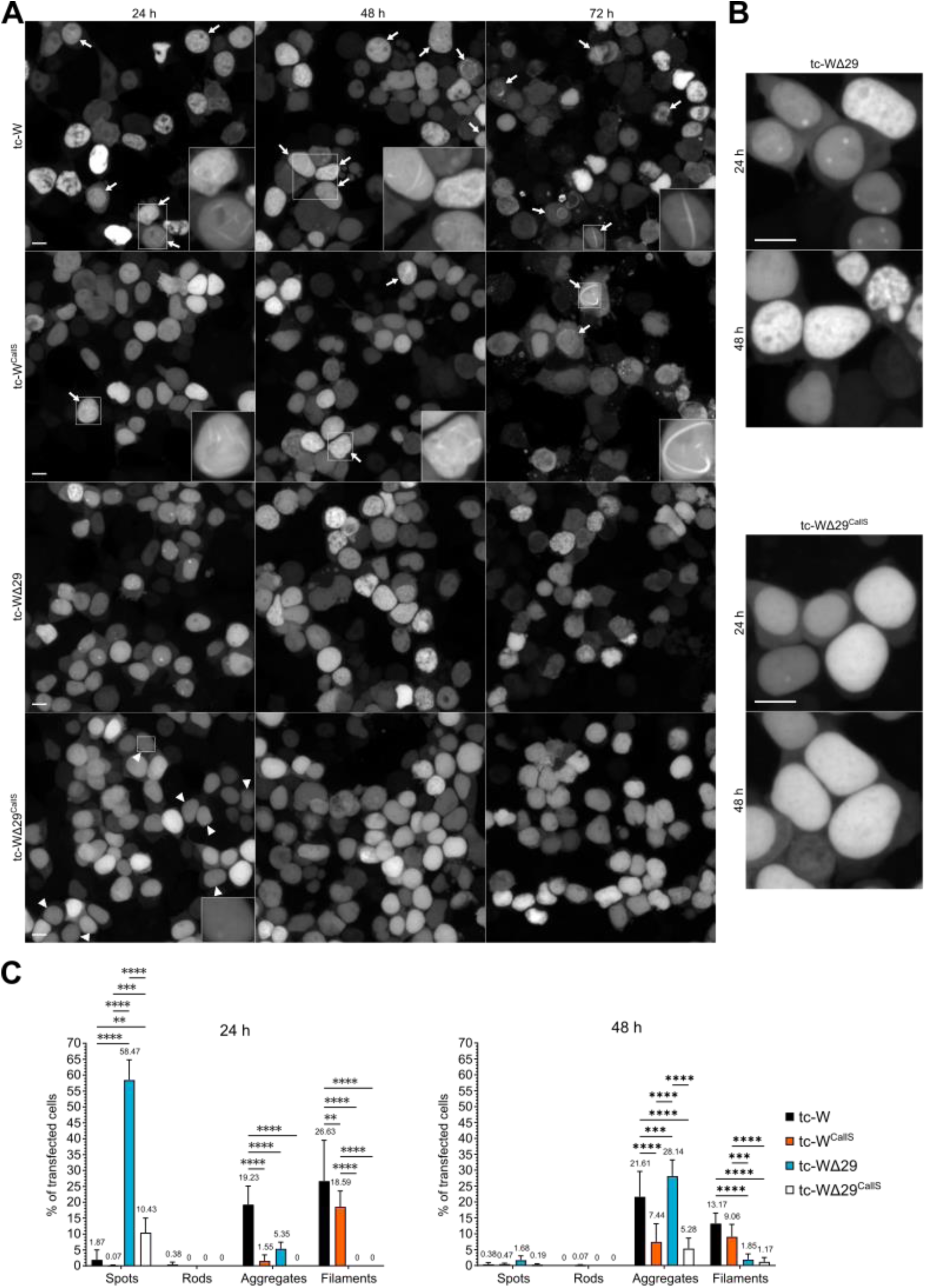
Δ29 deletion strongly impairs W^HeV^ ability to form filaments while CallS substitution impedes formation of non-filamentous condensates *in cellula*. (**A**) Z-stack projections of live HEK293T cells transfected with the indicated constructs, stained with ReAsH and imaged by confocal microscopy at 24h, 48h, and 72h post-transfection. Arrows indicate filament-containing nuclei, and arrowheads indicate nuclei with small, low-intensity spots for the WΔ29^CallS^ variant. Scale bars, 10 µm. (**B**) Closeup of cells expressing the indicated W^HeV^ variants, showing spots and amorphous aggregates for *tc*-WΔ29, and mostly nuclei with homogenous, diffuse signal for *tc*-WΔ29^CallS^. Scale bars, 10 µm. (**C**) Corresponding quantification of the percentage of transfected cells harboring the different nuclear structures formed by W^HeV^ at 24h and 48h post-transfection. Data are expressed as mean ± sd and analyzed by two-way ANOVA followed by a Tukey’s multiple comparisons test. \*\**p*<0.01, \*\*\**p*<0.001, \*\*\*\**p*<0.0001.

Replacing all the cysteine residues with serines (C^all^S variant) induces a significant decrease in the number of cells with nuclear filaments (**Figure 9A, C**). However, the most striking feature of this variant lies in its inability to form spots and rods and in its ability to form only very few irregular aggregates (**Figure 9A, C**). The WΔ29^CallS^ variant is both unable to form filaments and non-filamentous condensates (**Figure 9A-C**). Importantly, no apparent difference in expression level was observed among the various W^HeV^ variants (**Supplementary Figure S8**), thus ruling out the possibility that the observed differences could merely reflect differences in expression levels.

Altogether, these data indicate that W^HeV^ can give rise to filaments in cells and that this ability is impaired upon removal of the first 29 residues. The presence of cysteines – presumably oxidized (see below) – is not strictly required for the formation of filaments, as judged from the fact that the C^all^S variant is still able to form filamentous structures. Thus, while *in vitro* cysteines appear to be necessary, though not sufficient, for fibril formation, they appear dispensable for the formation of nuclear filaments. On the other hand, substituting cysteines with serines abrogates the ability of W^HeV^ to form spots, rods and, to a lesser extent, irregular aggregates, advocating for a scenario where the formation of these non-filamentous condensates would be mediated by disulfide bridge-driven dimers / oligomers.

### Both W expression and HeV infection induce reactive oxygen species production

Oxidative stress is a common feature during viral infections where mitochondria are targeted by viral proteins to promote a strong production of reactive oxygen species (ROS) (Foo et al., 2022), with this being also true in the case of NiV (Amurri et al., 2024; Escaffre et al., 2015). As such, henipavirus infection would generate an environment favorable for the formation of disulfide-bridged dimers/oligomers that mediate the formation of spots, rods and irregular aggregates, as judged from their absence / scarcity in cells transfected with the C^all^S construct (**Figure 9**). Since these condensates could be detected in the nuclei of cells transfected with *tc*-W, we expected transfection and/or expression of the W^HeV^ protein to mimic infection and to be able to trigger the production of ROS, thereby enabling ensuing aggregation. Quantification of ROS by confocal microscopy in HEK293T cells transfected with an empty vector or with a construct driving the expression of W^HeV^ confirmed that expression of the latter did significantly induce ROS production (**Figure 10A**, **B**). By contrast, transfection with the C^all^S construct did not trigger an increase in ROS (**Figure 10A**, **B**). Since the C^all^S variant is still able to form filaments, although to a lesser extent compared to the WT protein, but is severely impaired in its ability to form other types of condensates, results are consistent with a scenario where the “culprits” of the oxidative stress would be the latter rather than the filamentous structures. The increase in ROS triggered by the non-filamentous condensates would create an oxidative environment further favoring aggregation thereby amplifying the process. In line with expectations, experimental quantification of ROS upon infection, showed an increase in ROS in infected cells (**Figure 10C**).

**Figure 10.**
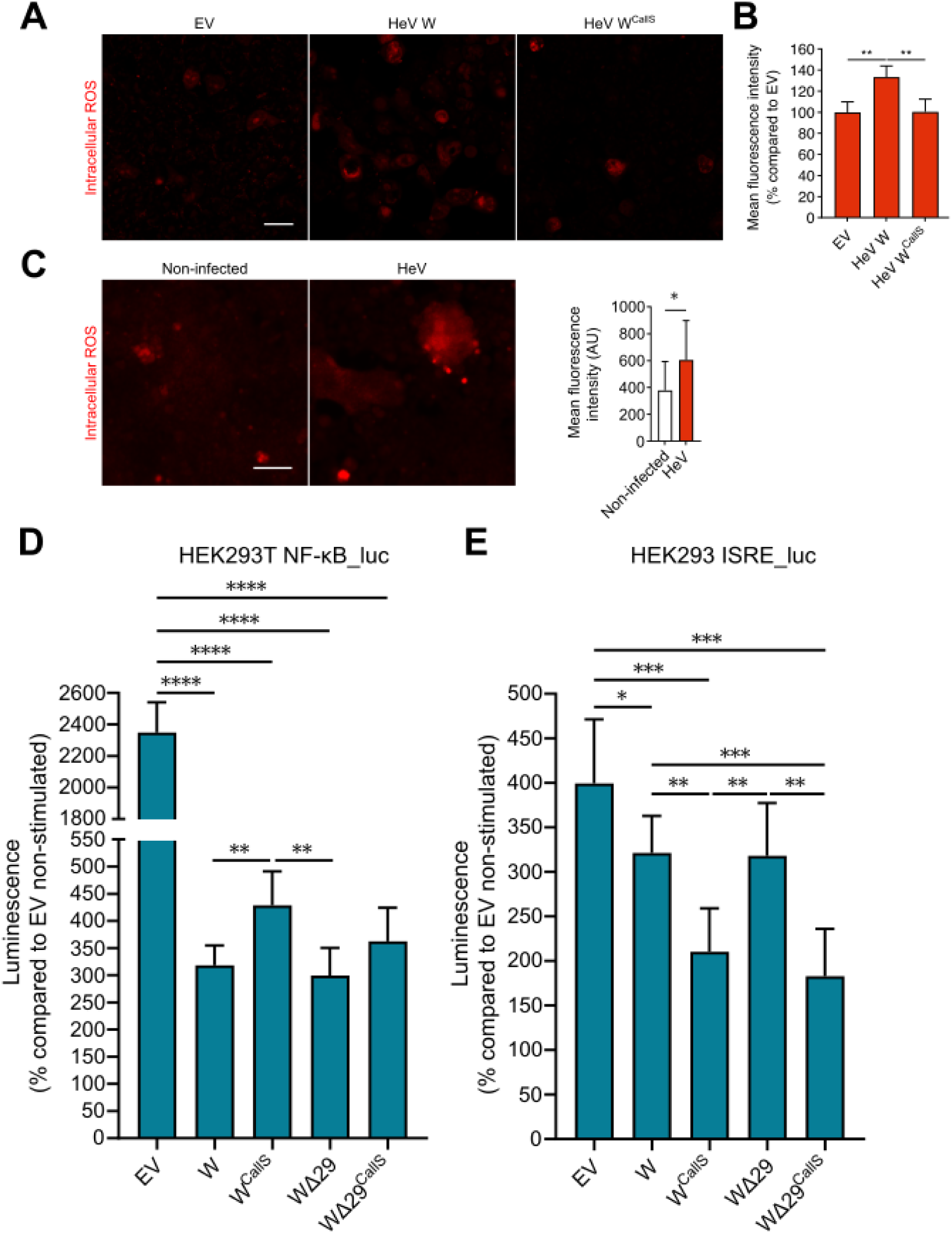
W^HeV^ condensation induces ROS production and impacts innate immunity pathways. (**A**) Detection of intracellular ROS in W^HeV^-transfected HEK293T cells at 24h post-transfection by confocal microscopy and (**B**) Fluorescence intensity quantification for each condition. Data are expressed as mean ± sd (n = 9 fields of view). Statistical significance was assessed with the Kruskal-Wallis test completed by a Dunn’s multiple comparison test. \*\**p* < 0.01. Scale bar, 20 µm. (**C**) Detection of intracellular ROS in HeV-infected HEK293T cells at 24h post-infection by epifluorescence microscopy and corresponding fluorescence intensity quantification for each condition. Data are expressed as mean ± sd (n = 9 fields of view). Statistical significance was assessed with the Mann-Whitney. \**p* < 0.05. Scale bar, 20 µm. (**D**) Measurement of NF-κB pathway activity in HEK293 NF-κB_luc reporter cells transfected with the indicated constructs. HEK293T NF-κB_luc cells were transfected and stimulated 24h later with IL-1β (50 ng/mL) before assessing NF-κB activity by luminescence quantification. Data are expressed as mean ± sd. Statistical significance was assessed with unpaired t tests. *p<0.05, ** p < 0.01, ***p < 0.001, ****p<0.0001. (**E**) Measurement of IFN response pathway activity in HEK293 ISRE_luc reporter cells transfected with the indicated constructs. HEK293 ISRE_luc cells were transfected and stimulated 24h later with IFN (100 U/mL) before assessing IFN pathway activity by luminescence quantification. Data are expressed as mean ± sd. Statistical significance was assessed with unpaired t tests. *p<0.05, ** p < 0.01, ***p < 0.001, ****p<0.0001

### W^HeV^-mediated repression of NF-κB pathway is impaired in the C^all^S variant that conversely supports enhanced repression of IFN signaling

After having shown that W^HeV^ forms various types of condensates in transfected cells, we sought at elucidating their functional impact. W proteins interact with multiple innate immune host factors to counteract the antiviral response (see (Gondelaud et al., 2022) and references therein cited). We recently showed that W^NiV^ inhibits NF-κB pathway activation *via* sequestration of 14-3-3 into the nucleus (Enchery et al., 2021). W proteins also counteract type I IFN (IFN-I) signaling by sequestering STAT1 into the nucleus (Quarleri et al., 2022; Shaw et al., 2004). We thus focused on the NF-κB and IFN-I pathways in our quest of the condensate functional impact.

To assess the possible impact on the NF-κB pathway, we used a HEK293T cell line stably expressing a firefly luciferase reporter under the control of a synthetic NF-κB promoter (HEK293T NF-κB_luc) (Pène et al., 2014). We transfected these cells with the W constructs driving the expression of the various W^HeV^ variants and measured luciferase activity as a proxy for NF-κB pathway activity. Luciferase expression, strongly induced by IL-1β treatment, was significantly inhibited in cells transfected with W^HeV^, while inhibition was found to be attenuated for the C^all^S variant (**Figure 10D**). These results mirror the propensity of these two W constructs to form condensates, with WT W^HeV^ being able to form spots, rods and irregular aggregates and C^all^S being unable to form spots and rods and less able to form irregular aggregates (**Figure 9**). By contrast, the WΔ29 variant, which is unable to form filaments, behaves like the WT W^HeV^ protein in terms of its ability to inhibit the NF-κB pathway (**Figure 10D**), arguing for a lack of impact of filaments on this pathway. The WΔ29^CallS^ variant tends to be less efficient for the repression of the pathway compared to the WT protein and the WΔ29 variant, supporting the critical role of the cysteines.

To assess the possible impact on the IFN response pathway, we used a HEK293 cell line stably expressing a firefly luciferase reporter under the control of a synthetic Interferon-Stimulated Response Elements (ISRE) (HEK293 ISRE_luc) (Lucas-Hourani et al., 2013). We transfected these cells with the same W constructs described above and measured luciferase activity as a proxy of IFN-I pathway activity (**Figure 10E**). Interestingly, the C^all^S variant inhibits the IFN-I pathway more efficiently than the WT W^HeV^ protein (**Figure 10E**) displaying therefore an opposite trend with respect to its impact on the NF-κB pathway (cf. **Figure 10D** and **10E**). Given the unique condensation properties of this variant, it is plausible that its enhanced ability to inhibit the IFN-I pathway reflects a scenario where it would be the monomeric form of W^HeV^ that would be responsible for STAT1 sequestration.

On the other hand, abrogating the ability to form filaments (WΔ29 variant) has no impact on the W-mediated inhibition of the IFN-I pathway, mirroring results obtained when analyzing the impact on the NF-κB pathway. Altogether, and as summarized in **Table 1**, these observations suggest that the W^HeV^-mediated inhibition of the NF-κB pathway is associated to the ability of W^HeV^ to form condensates whose formation is mediated by disulfide-bridged oligomers, and not merely on its presence as a monomeric form. By contrast, the latter condensates would be poorly able to sequester STAT1 and the only molecular species able to counteract IFN-I signaling would be the monomeric form of W^HeV^. In other words, while formation of spots, rods and aggregates, but not of filaments, is correlated to an enhanced W-mediated inhibition of the NF-κB pathway, favoring a primarily monomeric form of W (*via* the C^all^S substitution that impedes dimerization/nucleation) enhances the W-mediated inhibition of the IFN-I response pathway.

**Table 1.** Overview of the *in cellula* condensation properties and impact on innate immune pathways of the W^HeV^ variants.

| Properties<br>Variants | Formation of<br>filaments | Formation of<br>spots | Formation of<br>irregular<br>aggregates | Antagonism of<br>NF-κB<br>pathway | Antagonism of<br>IFN response<br>pathway |
| --- | --- | --- | --- | --- | --- |
| W | ++ | + | ++ | ++ | + |
| W<sup>C<sup>all</sup>S</sup> | + | - | + | + | ++ |
| WΔ29 | - | +++ | ++ | ++ | + |
| WΔ29<sup>C<sup>all</sup>S</sup> | - | ++ | +/- | + | ++ |

## Discussion

We herein report the crucial implication of cysteine residues in the fibrillation ability of the W protein *in vitro*. SEC analysis of W proteins preincubated for different periods under denaturing and oxidative conditions showed that they undergo disulfide bridge-mediated oligomerization. This process is closely related to W^HeV^ and W^NiV^ solubility since in the absence of disulfide-bridged oligomers, amorphous aggregates are formed, as shown by experiments in the presence of a reducing agent or using the W^HeV^ C^all^S variant. Oligomerization of both W^HeV^ and W^NiV^ proteins was also found to be closely tied to the formation of fibrils and as such, oligomers represent the fundamental drivers of primary nucleation. The requirement for a single cysteine residue for fibril formation and the substantial increase in W^HeV^ C^all^S solubility and ability to form fibrils upon generation of chemically-induced dimers support the conclusion that dimeric species are the primary drivers of nucleation. Notably, those dimers can be formed through any cysteine residue, as judged from the fibrillation abilities of the Cys123^only^, Cys316^only^, Cys334^only^ and Cys419^only^ variants. Furthermore, dimers do not require to be disulfide bridged to promote nucleation as illustrated by the fibrillation abilities of the artificially induced C^all^S dimers as obtained *via* addition of AP20187 to FKBP-C^all^S. Nucleation *via* disulfide-bridged dimers has been described in the case of Tau (Nizynski et al., 2017) and fish β-parvalbumin (Werner et al., 2020) while non-covalent dimers were shown to initiate nucleation of other fibrillogenic proteins, such as the Aβ peptide (Press-Sandler and Miller, 2019) or α-synuclein (Lan-Mark and Miller, 2022; Zhang et al., 2022). However, unlike these latter proteins, the monomeric species of W^HeV^ and W^NiV^ are not stable in solution and rapidly aggregate. W^HeV^ and W^NiV^ disulfide-bridged oligomers seem to enable bypassing the lag phase that is usually observed during the amyloidogenesis process (Alraawi et al., 2022) through the formation of covalent oligomers that eventually polymerize. Interestingly, mature W^HeV^ and W^NiV^ fibrils are not completely sensitive to reduction, indicating that a successive and distinct stabilizing mechanism, presumably brought by the PNT1 region (and in particular by the residues 2-29) (Gondelaud et al., 2025), is at play in the fibril elongation step compared to the initial nucleation step.

Another consequence of the presence of multiple Cys in W^HeV^ is the formation of intrachain disulfide bonds, which decreases the hydrodynamic radius of the monomer. This compact conformation is not required for W^HeV^ fibrillation, as shown by the Cys^only^ W^HeV^ variants. On the contrary, intrachain disulfide bonds may impair the fibrillation process by decreasing the nucleation extent since less cysteine residues are available to establish an interchain bridge. This scenario is in agreement with previous observations on other fibrillogenic proteins, where increasing the compaction of α-synuclein *via* the addition of an artificial disulfide bond strongly decreases its fibrillation propensity (Carija et al., 2019). Similarly, intrachain disulfide bonds in 4R Tau isoforms cause the formation of compact monomers that are not able to fibrillate (Nizynski et al., 2017). According to our observations, the probable function of cysteine residues is 1) to allow monomers to dimerize, enabling fibril nucleation, 2) to maintain them soluble and 3) to prevent fibril clustering.

Interestingly, using a tetracysteine tag already successfully employed in (Léger et al., 2020), we showed that *in cellula* W^HeV^ forms different types of condensates. It should be noted that although LLPS is often discussed synonymously with biomolecular condensation, the latter is in fact more general, covering processes beyond just liquid–liquid demixing and including gelation, crystallization, polymerization, clustering, and amorphous or amyloid aggregation (Alberti et al., 2019). W^HeV^ was found to form giant filamentous structures with an exclusive nuclear localization. In light of the differences in size, filaments cannot be regarded as the direct cellular counterpart of fibrils observed *in vitro.* However, one can speculate that giant filaments are composed of bundles of fibrils, a hypothesis that would need to be ascertained using high-resolution ultrastructural analysis, as described for the fibrillar condensates formed by the NSs protein from the Rift Valley fever virus (RVFV) (Léger et al., 2020). In addition, the WΔ29 variant is unable to form filaments, a finding that parallels the failure of this variant to fibrillate *in vitro* thereby supporting the hypothesis of a close relationship between filaments and fibrils.

Fibril formation *in vitro* appears to take place through the *deposition* pathway rather than through the *condensation and maturation* pathway (Vendruscolo and Fuxreiter, 2022), as judged from the lack of evidence in support of a droplet state preceding the formation of fibrils. Mirroring these findings, filaments observed *in cellula* do not appear to result from *maturation* of globular condensates. Incidentally, the latter were found to have a seemingly solid-like nature. Whether globular condensates contain fibrils of the same length as those observed *in vitro* will be investigated in future studies.

Substitution of cysteine residues with serines (C^all^S variant) impairs formation of non-filamentous condensates (spots, rods and irregular aggregates), whose formation therefore appears to be mediated by disulfide bridge-driven dimers / oligomers. The reliance on oxidized cysteines for fibril formation *in vitro* raises the question of whether W proteins can form condensates mediated by disulfide bridges *in cellula*, given the reducing environment of the nucleus (and of the cytosol). Noteworthy, NiV infection induces ROS production (Escaffre et al., 2015) as does HeV infection (**Figure 10C**), and we showed here that the sole ectopic expression of W^HeV^ resulted in the production of ROS, thereby creating an environment favorable to the formation of condensates relying on the formation of disulfide bridges and eventually provoking and/or amplifying W^HeV^ condensation. These observations are in line with several studies on various proteins whose condensation in fibrillar structures relies on disulfide bridges. The human protein p16^INK4A^ that only fibrillates when its unique cysteine is oxidized, was shown to fibrillate in cultured cells under mild oxidizing conditions (Göbl et al., 2020). Likewise, disulfide bond-dependent viral amyloids made by the RVFV NSs protein could be observed in both transfected and infected cells (Léger et al., 2020). These two examples, together with the ones presented herein, indicate that the oxidation of cysteine residues shown to be critical for protein condensation does take place in the cellular context.

In NiV-infected cells, equimolar proportions of P, V, and W transcripts are detected in the early stage of infection while the quantity of V and W transcripts increases as the infection progresses (Shaw, 2009). V and W proteins are therefore synthesized at an early stage after infection. The quick formation of W condensates may benefit to the virus to fight against the cell innate immune response in light of the short timescale of viral infections, the duration of which is limited to few hours *in vitro* (Marsh et al., 2013) and few days *in vivo* after experimental infection (Guillaume et al., 2009). As previously described (Enchery et al., 2021), W^NiV^ CTD binds and accumulates 14-3-3 into the nucleus leading to an increased cytosolic translocation of NF-κB p65/p50 and consequently a reduced expression of proinflammatory cytokines. Our study confirms and extends these findings: not only it shows that the W^HeV^ protein is able to impair the NF-κB pathway, but it also sheds light on the molecular underpinnings. Indeed, the formation of spots, rods and irregular condensates was shown to augment this ability, as judged from the observation that the NF-κB pathway is less impaired in the C^all^S variant that is unable to form this type of condensates *in cellula*. By contrast, it is the monomeric form of W^HeV^ that appears to be more efficient to counteract IFN-I signaling. As such, the virus would need both non-assembled and self-assembled forms of W to ensure efficient blockade of the host’s innate immune response. Ongoing and future studies will address STAT1 co-localization with the various types of W^HeV^ condensates.

Contrary to RVFV, whose disulfide-bridge mediated NSs large filaments were shown to be associated with IFN-I response silencing (Léger et al., 2020), the functional impact of W^HeV^ filaments remains elusive and deserves future investigation. The herein uncovered impact of W^HeV^ redox-dependent, non-filamentous condensates on the NF-κB pathway, aligns with previous reports pointing out the critical role of viral condensates in inhibition of the antiviral response (refer to (Cai et al., 2023) as an example) while further broadening the biological relevance of biocondensates in host immune evasion to encompass non-liquid forms.

In conclusion, the present study constitutes the first experimental evidence of the ability of the *Henipavirus* W proteins to self-assemble in a cellular context. By establishing a functional link between redox-dependent condensates and impairment of the NF-κB pathway, this study provides the first clues on the functional impact of these latter condensates. In addition, by showing a link between W oligomerization and reduced ability to inhibit the IFN-I response, our study provides hints on the mechanisms underlying STAT1 sequestration, a key determinant of *Henipavirus* pathogenicity. Our study thus contributes to shed light on the possible molecular mechanisms by which this virulence factor may contribute to *Henipavirus* pathogenesis. Further studies will be necessary to explore to which extent W condensation participates to W function and potentially to the switch from inhibition of the IFN-I pathway to inhibition of NF-κB signaling in the context of *Henipavirus* infection. The present study sets the stage for future investigations aimed at assessing the generality of this mechanism in other paramyxoviruses expressing a P-encoded W protein. From a more general perspective, it also adds to the hitherto relatively few examples of viral fibrillar and non-fibrillar functional condensates and thus broadens our knowledge on their use by viruses as a strategy to counteract host cell functions.

## Acknowledgments

We acknowledge the PICsL-FBI electron microscopy facility of the IBDM, members of the national infrastructure France-BioImaging supported by the French National Research Agency (ANR-10-INBS-04), for providing access and thank Nicolas Brouilly and Fabrice Richard for their technical help and guidance. We also acknowledge Patrick Fourquet from the mass spectrometry facility of Marseille Proteomics (marseille-proteomique.univ-amu.fr), supported by IBISA (Infrastructures Biologie Santé et Agronomie), Plateforme Technologique Aix-Marseille, the Cancéropôle PACA, Région Sud-Alpes-Côte d’Azur, the Institut Paoli-Calmettes, the Centre de Recherche en Cancérologie de Marseille (CRCM), Fonds Européen de Développement Régional and Plan Cancer for mass spectrometry analyses. We thank María Maté for the SEC-MALLS experiments. We thank all the AFMB technical and support staff (Denis Patrat, Patricia Clamecy, Béatrice Rolland, Paul Zamboni, Chantal Falaschi and Fabienne Amalfitano). We also thank Joanna Brunel who built the parental plasmid encoding FKBP from synthetic gene constructs, Louis-Marie Bloyet for sharing the pCAGGS vector and Pierre-Olivier Vidalain (all from CIRI, Lyon) for providing the BEAS-2B and HEK293 ISRE_luc cells, and finally Racha Majed (Refeyn company) for the mass photometry measurements. We acknowledge the contribution of SFR Biosciences (UAR3444/CNRS, US8/Inserm, ENS de Lyon, UCBL) facility Plateau Technique d’Imagerie/Microcopie (PLATIM). We acknowledge the team of the INSERM Jean Mérieux BSL-4 lab (Lyon).

## Data Availability

The data presented in the current study are available from the corresponding authors upon reasonable request.

## Declaration of generative AI and AI-assisted technologies in the writing process

During the preparation of this work the author(s) used ChatGPT in order to polish the style of a few sentences throughout the text. After using this tool/service, the author(s) reviewed and edited the content as needed and take(s) full responsibility for the content of the published article.

## Materials and Methods

### Generation of constructs

The generation of the constructs is detailed in the **SI Appendix, Materials and Methods** (see **Supplementary Tables S3** and S**4**). The resulting constructs for bacterial and eukaryotic expression allow the expression of 6-His tagged recombinant proteins, and of N- or C-terminally *tc*-tagged proteins, respectively.

### Expression and purification of recombinant proteins

Expression of all recombinant proteins was made in the *E. coli* strain T7pRos. The detailed protocol for protein expression and purification is detailed in the **SI Appendix, Materials and Methods**. Briefly, recombinant W proteins and their variants were purified in denaturing and reductive conditions (*i.e.* in the presence of 6 M urea and of 10 mM DTT) by Ni-NTA affinity chromatography and anion exchange chromatography (AEC). Protein samples were then desalted to remove the DTT and stored at -80°C in denaturing conditions. FKBP-C^all^S was purified in the same way except that an additional preparative SEC was performed in HBS/1 M urea. Purified proteins analyzed by SDS-PAGE are shown in **Supplementary Figure S9**.

### Protein sequencing by tandem mass spectrometry

The identity of the compact conformation of W^HeV^ was confirmed by mass spectrometry analysis of tryptic fragments obtained after digestion of the purified protein band excised from SDS-polyacrylamide gel. The detailed protocol is further described in **SI Appendix, Materials and Methods**.

### Analytical SEC of non-incubated and incubated proteins

One hundred µL of W^HeV^ (WT and variants) or W^NiV^ (WT) samples at 75 µM in HBS/6 M urea buffer, incubated at 37°C without agitation for different times, were injected onto a Superdex S200 Increase 10/300 GL (Cytiva) column equilibrated in HBS and eluted in the same buffer at a flow rate of 0.8 mL/min and at room temperature.

### Mass photometry measurements

Mass photometry experiments were carried out on a TwoMP mass photometer (Refeyn). WT W^HeV^ was incubated at a final concentration of 75 µM for 48h at 37°C and then diluted to 100 nM in HBS for molecular mass measurements.

### Turbidimetry and fluorimetry measurements

Protein samples were incubated at 37°C for different times (indicated in each experiment) as detailed in the **SI Appendix, Materials and Methods**. Protein solutions were then buffer exchanged to remove urea. The resulting urea-free protein samples were analyzed in the presence of thioflavin T (ThT). Turbidity measurements and fluorescence measurements were performed on a Tecan microplate reader GENios Plus in black 96-well plates with transparent flat bottom (Greiner, 655096) incubated at 37°C.

The same protocol was followed in the experiments made in the presence of DTT. 2 mM DTT was added either just after the buffer exchange step (t_0_), or on a W^HeV^ sample that had already been incubated for 2h (t_2h_). In this latter case, turbidity and fluorescence were monitored for two extra hours (t_2h + 2h_). This latter sample was further incubated at 37°C for a total of 24h and then centrifuged at 10 000 x *g* for 10 min. The pellet was resuspended in 150 µL of HBS before being deposited on a NS-EM grid.

Measurements were performed at least in triplicate (indicated in each experiment) and data are represented as mean ± standard deviation (s.d.). The statistical analysis used is detailed for each experiment, using either a one-way ANOVA with the Dunnett’s multiple comparison test, or a Student’s t-test.

### Negative-staining electron microscopy (NS-EM)

NS-EM carbon coated grids were prepared from protein samples incubated for 120 min at 37°C in HBS. A final concentration of 2 µM was used for this purpose as detailed in the **SI Appendix, Materials and Methods**. Measurements of fibrils lengths were made manually with the ImageJ software (Schneider et al., 2012).

### Chemically-induced dimerization of FKBP-C^all^S

FKBP-C^all^S sample was incubated in the absence or presence of AP20187 (Sigma-Aldrich, SML2838) as detailed in the **SI Appendix, Materials and Methods**. A part of the sample was analyzed by SEC and excess AP20187 was removed using a PD Minitrap G-25 column (Cytiva) equilibrated in HBS. Buffer exchanged samples were analyzed by turbidity and fluorimetry and NS-EM in the same way as described above.

### Multi-angle laser-light scattering (SEC-MALLS)

The formation of FKBP-C^all^S dimeric species upon addition of the AP20187 molecule was also assessed by SEC-MALLS. For this purpose, the same protocol as previously described was followed and the sample was centrifuged for 15 min at 10 000 x *g* at room temperature. 50 µL were injected at a flow rate of 0.6 mL/min on a Superdex S200 Increase 10/300 GL (Cytiva) column equilibrated in HBS controlled by an Ultimate 3000 HPLC (Thermo Scientific). Measurements were carried out on a Wyatt Dawn 8 angles MALLS equipped with an Optilab refractometer (Wyatt).

### W^HeV^ C^all^S variant crosslinking by DTSSP

W^HeV^ C^all^S was cross-linked in the presence of 2 mM DTSSP (Pierce, 21578). The entire protocol is described in the **SI Appendix, Materials and Methods**. The sample was analyzed by SEC and by turbidimetry and fluorimetry after a buffer-exchange step in HBS, as described above, at a final concentration of 30 µM. NS-EM grids were prepared after 2h of incubation in HBS.

### Cellular cultures

Human embryonic kidney 293T (HEK293T), A549 and Vero E6 cells were cultured in Dulbecco’s Modified Eagle Medium (DMEM) (Gibco, Cat# 61965-026) supplemented with 10 % fetal bovine serum (FBS) (DMEM/10% FBS). The HEK293T cell line stably transduced with a firefly luciferase reporter under the control of a synthetic NF-κB promoter (HEK293T NF-κB_luc), and the HEK293T cell line expressing an ISRE-dependent luciferase reporter (HEK293 ISRE_luc, kindly provided by the team of Pierre-Olivier Vidalain) were cultured in DMEM/10% FBS, with 200 μg/mL hygromycine B (Invitrogen, Cat# 10687010) or 500 μg/mL G-418 (MedChemTronica, Cat# HY-17561), respectively. BEAS-2B cells (kindly provided by the team of Pierre-Olivier Vidalain) were cultured in DMEM/F-12 (Gibco, Cat# 31331-028) with 10% FBS. All the cells were cultured at 37 °C in a 5 % CO_2_ incubator.

### Virus

WT HeV (Australia/1994/Horse18 isolate, GenBank accession number MN062017.1) was initially isolated from a horse and obtained from Porton Down Laboratory, UK (Dhondt et al., 2013; Guillaume et al., 2009; Guillaume-Vasselin et al., 2016). Briefly, viral stock was produced by infecting Vero E6 cells at a multiplicity of infection of 0.001 in Opti-MEM (Gibco, Cat# 51985042). After 1 h incubation at 37°C, the medium was replaced with DMEM with 5% FBS and cells were incubated at 37°C at 5% CO_2_ for 48h. Viral supernatants were collected and centrifuged (400*g*, 10 min, 4°C), aliquoted and titrated in plaque forming units by classic dilution limit assay on Vero E6 cells.

### Immunofluorescence

Transfected HEK293T cells were stained using standard immunofluorescence protocol detailed in the **SI Appendix, Materials and Methods**, using a rabbit anti-W^CTD^ antibody (GeneScript) (1:100) and Alexa Fluor 647 Donkey anti-Rabbit IgG (H+L) Highly Cross-Adsorbed Secondary antibody (Invitrogen, Cat# A-31573) (1:500), before imaging on a Yokogawa HCS CQ1 spinning disk confocal system (PLATIM, SFR Biosciences Lyon).

### Live imaging and quantification of W^HeV^ condensates

HEK293T, BEAS-2B and A549 cells were seeded in Ibidi µ-Slide 8 Well and transfected 24h later with 0.4 µg of the indicated constructs, using FuGENE 6 with a DNA:transfection reagent ratio of 1:4, or TransIT-LT1 for A549 cells with a 1:3 ratio. They were stained at the indicated times post-transfection using the TC-ReAsH or FlAsH II In-Cell Tetracysteine Tag Detection Kit and 1 µg/mL Hoechst 33342 when stated. To this end, cells were rinsed with Opti-MEM, incubated for 45 min at room temperature with 2.5 µM of ReAsH/FlAsH-EDT2 labeling reagents in Opti-MEM, and finally rinsed with Opti-MEM, allowing the visualization of *tc*-tagged W^HeV^ on a Yokogawa HCS CQ1 confocal system. Z-stacks of 10 to 15-µm-range with an imaging step of 0.7-1 µm were acquired in random areas of the wells. Maximum intensity projections were generated automatically by the microscope software. Quantification of the percentage of cells bearing condensates was performed with Fiji (v2.9.0). The total number of stained cells on the Z-projections were counted semi-automatically using Threshold, Fill Holes, Watershed and Analyze Particles functions with constant parameters between projections (minimum size of 3000 px^2^). Cells with condensates were counted manually.

### Time-lapse imaging

HEK293T cells were seeded in Ibidi µ-Slide 8 Well (175,000 cells/well) previously coated with 50 µg/mL poly-D-lysine. Cells were transfected 24h later with 0.4 µg of the pCAGGS plasmid encoding tc-W with FuGENE 6, with a DNA:transfection reagent ratio of 1:4. Different staining and imaging conditions using a Yokogawa HCS CQ1 confocal system (40x objective UPLSAPO40X2, Olympus) were tested. For Figure 6A, 6C, and S7B, cells were rinsed 24h later with Opti-MEM and 300 µL of Opti-MEM with 1.5% FBS and 0.33 µM FlAsH were added. After 30 min of incubation at room temperature, cells were imaged every 50 min with a 488 nm laser (25% power) and 2.5 s exposition, at 37°C and with 5% CO_2_. For Figure 6B, cells were rinsed 24h later with Opti-MEM and 300 µL of Opti-MEM with 0.5% FBS and 0.67 µM FlAsH were added. After 30 min of incubation at room temperature, cells were imaged every 15 min with a 488 nm laser (1.5% power) and 3 s exposition, at 37°C and with 5% CO_2_. For Figure 7E, cells were rinsed 24h later with Opti-MEM and 300 µL of Opti-MEM with 2 µM FlAsH were added. After 30 min of incubation at room temperature, cells were imaged every 60 min with a 488 nm laser (7% power) and 2.5 s exposition, at 37°C and with 5% CO_2_. For Figure S7A, cells were rinsed 24h later with Opti-MEM and 300 µL of Opti-MEM with 1.5% FBS and 2 µM FlAsH were added. After 30 min of incubation at room temperature, cells were imaged every 60 min with a 488 nm laser (7% power) and 2.5 s exposition, at 37°C and with 5% CO_2_. For Figure S7C, cells were rinsed 24h later with Opti-MEM and 300 µL of Opti-MEM with 0.67 µM FlAsH were added. After 30 min of incubation at room temperature, cells were imaged every 15 min with a 488 nm laser (1.5% power) and 3 s exposition, at 37°C and with 5% CO_2_.

### High-resolution confocal microscopy of condensates

HEK293T cells were seeded in Ibidi µ-Slide 8 Well and transfected as previously described with FuGENE 6. At 72h post-transfection, cells were rinsed with Opti-MEM, incubated for 45 min at room temperature with 2.5 µM ReAsH in Opti-MEM, and finally rinsed with Opti-MEM. They were next imaged in Opti-MEM on an abberior Instruments INFINITY Line confocal microscope with a 60x oil immersion objective (UPLXAPO60XO, NA 1.42, Olympus). To improve resolution, reduce background noise and more globally increase the signal-to-noise ratio in order to be able to segment and quantify the condensates, the MATRIX detector was used for the acquisitions, requiring a pinhole size >10, and a pixel size of 65 nm. The following parameters were used for the acquisitions post-treatment: Differential detection between 0.8 and 1.5 and Sharpening between 6 and 11. Analysis of the sphericity of the spots were performed with Fiji by cropping the stacks to keep only the individual spots, a Gaussian blur of 4 was then applied, images were thresholded, and the sphericity and volumes were finally measured with MorphoLibJ plugin (Legland et al., 2016) using the Analyze Regions 3D function.

### 3D Reconstruction and quantification of filamentous and round-shaped condensates

Z-stack images acquired by confocal fluorescence microscopy in multiple channels were imported into Imaris software (Bitplane, version 10.0) and converted to the “.ims” format for subsequent analysis. Filamentous structures were reconstructed using the “Volume” creation tool within the 3D View module. For each fluorescence channel, the “Add New Surfaces” function was used to initiate object segmentation. The “Classify Surfaces” and “Object-Object Statistics” features were enabled to allow detailed quantification of individual structures and their spatial relationships. To restrict the analysis to biologically relevant areas, the “Segment Only a Region of Interest” option was applied. Background fluorescence was reduced using the “Background Subtraction (Local Contrast)” algorithm. Surface boundaries were manually adjusted to accurately match the fluorescent signal observed in the original image slices. Additional filtering criteria, including “Quality,” “Mean Intensity,” “Sphericity,” and “Shortest Distance to Neighboring Object,” were applied to minimize artifacts or non-specific signals. The same thresholding parameters were then uniformly applied across all images within each experimental group to ensure consistency. Round-shaped condensates were analyzed following the same workflow, with the exception that filtering was limited to “Mean Intensity” and “Sphericity”. The “Smooth” option was enabled to refine surface contours, while the “Split Connected Items” option was disabled to preserve the continuity of individual filaments. After surface reconstruction, objects were categorized into subgroups based on parameters such as length, volume, and surface area. Quantitative data were exported for downstream statistical analysis. Subgroups were visually distinguished by adjusting surface color and transparency within the software.

### Preparation of samples for electron microscopy of transfected cells

HEK293T cells were seeded in 35-mm gridded dishes (Ibidi, Cat# 81166) at a density of 600,000 cells per dish. Cells were transfected 24h later as previously described using the TransIT-LT1 reagent with 3.2 μg of plasmid DNA (empty pcDNA3.1(+) vector or tc-tagged W construct). Live-cell imaging was performed 24h and 48h later using the TC-ReAsH™ II In-Cell Tetracysteine Tag Detection Kit on a Yokogawa HCS CQ1 confocal system to locate filament-bearing cells. ReAsH-EDT2 labeling reagent was used at a final concentration of 0.625 μM. Cells were imaged at 4 and 10x magnifications and Z-stacks of 8 to 12-μm-range with an imaging step of 0.8 μm were acquired with the 40x objective. Maximum intensity projections were generated automatically by the microscope software. After imaging, cells were fixed with 4% methanol-free formaldehyde (Electron Microscopy Sciences, Cat# 15713) for 10 min, and kept in DPBS before tomography analysis.

### Embedding and ultramicrotomy of transfected cells, and EM analyses

HEK293T cells were prepared following a regular EM sample preparation detailed in the **SI Appendix, Materials and Methods**. Briefly, cells were post-fixed in 1% OsO_4_, *en bloc* contrasted in 1% uranyl acetate, dehydrated in ethanol and embedded in Epon. The region of interest was recovered from the dish and targeted using Ibidi landmarks. One-micron thick sections were collected on a glass slide. Sections of interest (those corresponding to the relevant z-positions in the fluorescence stacks) were identified by toluidine blue staining and re-embedded in pure Epon 812 and incubated at 60°C for approximately 12h. The samples were then immersed in liquid nitrogen to remove the resin-embedded sections from the slide (cracking). 70-90 nm ultrathin sections were post-stained with uranyl acetate and lead citrate. Images were taken using a Tecnai™ T12 Spirit 120 kV.

### Assessment of cell death by fluorescence microscopy

HEK293T cells were seeded in Ibidi µ-Slide 8 Well (150,000 cells/well) previously coated with 50 µg/mL poly-D-lysine. Cells were transfected 24h later with 0.4 µg of the indicated plasmids with FuGENE 6, with a DNA:transfection reagent ratio of 1:4. The next day, activated Caspase-1 was stained using the Caspase 1 Staining Kit (Far Red) (Abcam, Cat ab270784). The staining solution was prepared according to the manufacturer’s instructions, and after removing the medium, 100 µL were added per well. Cells were incubated at 37°C for 45 min, then the staining solution was removed and the cells were rinsed twice with Opti-MEM. Cells were next stained with a mix of 2.5 µM FlAsH from the TC-FlAsH™ II In-Cell Tetracysteine Tag Detection Kit (Invitrogen, Cat T34561), 3 µg/mL propidium iodide, and 1 µg/mL Hoechst in Opti-MEM, at room temperature for 45 min in the dark. Cells were rinsed with Opti-MEM and imaged with a Yokogawa HCS CQ1 confocal system in Opti-MEM. Positive control cells were incubated with 0.01% Triton X-100 in Opti-MEM for 10 min prior to staining.

### Cell viability/metabolic assay

HEK293T cells were seeded in poly-D-lysine-coated 96-well black plate (Corning, 354640) and transfected 24h later with 0.2 μg of the indicated plasmids with the TransIT-LT1 reagent. Cell viability was measured the next day using alamarBlue Cell Viability Reagent (Invitrogen, DAL1025) by quantifying fluorescence intensity on a Tristar 5 microplate reader, according to the manufacturer’s instructions, using an excitation wavelength of 570 ± 7.5 nm, an emission wavelength of 585 ± 7.5 nm, and an exposure time of 100 ms.

### Luciferase assay

HEK293T NF-κB_luc and HEK293 ISRE_luc cells were seeded in 96-well plates (50,000 cells/well) previously coated with 50 µg/mL poly-D-lysine, and transfected 24h later with 0.2 µg of an empty pCAGGS vector or pCAGGS plasmids coding for *tc*-tagged W^HeV^ variants, using FuGENE 6 (Promega, Cat# E2691) according to the manufacturer’s instructions, with a DNA:reagent ratio of 1:4. The medium was replaced 24h later with DMEM/10% FBS supplemented or not with 50 ng/mL human IL-1β (PeproTech, Cat 200-01B) or 100 U/mL universal type I IFN (PBL Assay Science, Cat 11200-1). Luciferase activity was measured after 5 h of incubation at 37°C using the Bright-Glo Luciferase Assay System (Promega, E2620). The medium was replaced with 50 μL per well of DMEM/10 % FBS, and 50 μL per well of Bright-Glo Luciferase Assay Substrate reconstituted in assay buffer were added before quantifying luminescence in a 96-well white opaque plate (1 s per well) on a Tristar 5 microplate reader (Berthold Technologies GmbH & co.KG)

### Intracellular ROS quantification

Intracellular ROS were quantified in HEK293T cells seeded and transfected as described for immunofluorescence staining, using the Fluorometric Intracellular ROS Kit (Sigma-Aldrich, MAK145). Briefly, medium was replaced with DMEM/10% FBS and the ROS detection master mix was then added with a medium:master mix ratio of 0.9:1. Cells were incubated for 1 h in a 5% CO2, 37 °C incubator, and then imaged on a Yokogawa HCS CQ1 confocal microscope. For each condition, single optical sections from nine random areas in the wells were acquired. Images were then analyzed with Fiji (v2.9.0), by first subtracting the background (rolling ball radius of 50 pixels) and then measuring the mean gray value for each field of view. For ROS detection in infected cells, HEK293T cells seeded in Ibidi µ-Slide 8 Well were infected with 100 PFU of HeV in Opti-MEM or left uninfected. After 3 h of incubation at 37°C, the medium was replaced by DMEM/10% FBS, and ROS were quantified 24h later using the Fluorometric Intracellular ROS Kit. Cells were imaged with a Leica DMIRB epifluorescence microscope with a 10x objective and analyzed with Fiji.

### Western blot

HEK293T cells were seeded in 12-well plates (700,000 cells/well) and transfected 24h later with 1.4 µg of the indicated constructs using FuGENE 6 as previously described. The next day, cells were rinsed with ice-cold DPBS, collected on ice in RIPA buffer (Invitrogen, Cat 89901) supplemented with Halt Protease and Phosphatase Inhibitor Cocktail (Invitrogen, Cat 78442), and sonicated using a Bioruptor Plus (Hologic Diagenode). Protein samples were then prepared for gel loading in NuPAGE LDS Sample Buffer (Invitrogen, Cat NP0008) and NuPAGE Sample Reducing Agent (Invitrogen, NP0009), heated at 96°C for 10 min, and separated by electrophoresis using 10% Mini-PROTEAN® TGX™ Precast Protein Gels (Bio-Rad, Cat 4561036) in TG-SDS buffer (Euromedex, Cat EU0510). Separated proteins were then transferred on PVDF membranes using Trans-Blot Turbo Midi 0.2 µm PVDF Transfer Packs (Bio-Rad, Cat 1704157) and the Trans-Blot Turbo Transfer System (Bio-Rad, Cat 1704150). The membranes were then blocked with TBS buffer + 0.05% Tween 20 (TBST) + 5% milk for 1 h under agitation, rinsed with TBST and incubated with a rabbit anti-W^CTD^ antibody (GeneScript) or a mouse anti-GAPDH antibody (Sigma-Aldrich, Cat MAB374) diluted in TBST + 0.2% milk (both 1/1000), for 1 h at room temperature or overnight at 4°C, respectively. They were rinsed with TBST three times for 10 min, and incubated with Anti-Rabbit IgG (H+L), HRP Conjugate (Promega, Cat W4011) or Anti-Mouse IgG (H+L), HRP Conjugate (Promega, cat W4021) secondary antibodies diluted in TBST + 5% milk (both 1/10,000). After three rinses with TBST for 10 min, revelation was performed using the SuperSignal West Pico PLUS Chemiluminescent Substrate (Thermo Scientific, Cat 34580) on a ChemiDoc (Bio-Rad).

