## Supplemental text, figures and tables for "Cysteine Redox State Governs the Condensation Pathway of Hendra Virus W Protein and Differentially Impacts Type I IFN and NF-κB signaling"

#### Supplementary Materials and Methods

##### Generation of constructs

All the constructs for the bacterial expression of the W proteins were obtained by overlap PCR extension following the same protocol. PCRs 1 and 2 used the templates and primers (Eurofins Genomics) listed in **Supplementary Tables S3** and **S4**. After DpnI treatment, aliquots of PCRs 1 and 2 were mixed and used as template for PCR 3 along with primers attl1a and attl2a (Gruet et al., 2016). In some cases, PCR products were gel-purified. In all cases, PCR 3 product was inserted in Gateway plasmid pDEST17O/I via LR reaction (ThermoFisher). The pDEST17O/I expression vector is a modified pDEST17 vector (Invitrogen) in which the LacO and LacI encoding sequences were inserted upstream the T7 promoter in order to allow a better control of protein expression (Vincentelli et al., 2004). The resulting constructs allow the expression under the T7 promoter of N-terminally hexahistidine tagged recombinant proteins in which the native W sequence is preceded by a vector-encoded MSYYHHHHHHLESTSLYKKAGS amino acid stretch and by a TEV cleavage site (ENLYFQG).

The pcDNA3.1(+) $W^{HeV}$-*tc* construct driving the eukaryotic expression of $W^{HeV}$ with a C-terminal tetracysteine tag (*tc*) (FLNCCPGCCMEP) was obtained by PCR using Q5 DNA polymerase and GC-rich template enhancer, using the $W^{HeV}$ constructs in pDEST17O/I as template and primers indicated in **Supplementary Tables S3** and **S4**. After DpnI treatment, the resulting amplicon was digested with HindIII and XhoI and then ligated with HindIII/XhoI-digested pcDNA3.1(+). The pcDNA3.1(+) construct encoding

$W^{HeV}$ with an N-terminal *tc* (referred to as *tc*-$W^{HeV}$) was obtained using In Fusion HD Cloning technology (Takara, Cat# 639650). The amplicon coding for *tc*-$W^{HeV}$ was generated by PCR with PrimeSTAR GXL DNA Polymerase (Takara, Cat# R050A), using the $W^{HeV}$ construct in pDEST17O/I as template and primers indicated in **Supplementary Tables S3** and **S4**. The amplicon was gel-purified with NucleoSpin Gel and PCR Clean-up kit (Macherey-Nagel, Cat# 740609.250) and cloned into a HindIII/XhoI-digested and gel-purified pcDNA3.1(+) vector following In Fusion kit's instructions. The pCAGGS constructs driving the eukaryotic expression of either tc-tagged $W^{HeV}$ variants or mStayGold were generated similarly by cloning the corresponding amplicons or synthetic DNA fragments into XhoI-digested and gel-purified pCAGGS vector, as indicated in **Supplementary Tables S3** and S**4**, following instructions of the In-Fusion Snap Assembly cloning kit (TaKaRa, Cat 638949). The pCAGGS-tc-mStayGold plasmid encodes the green fluorescent protein mStayGold (Ando et al., 2024) fused to a N-terminal tc tag as a negative control of protein condensation.
All the constructs were checked by DNA sequencing (GeneWiz/Azenta and Eurofins Genomics) and found to conform to expectations.

**Expression and purification of recombinant proteins**

Expression of all recombinant proteins was made in the *E. coli* strain T7pRos. Cultures were grown overnight to saturation in LB medium containing 100 µg/mL ampicillin. The overnight culture was diluted 1/5 into 1 L of Turbo Broth medium supplemented with ampicillin and grown at 37°C for 3h, at 200 rpm before induction of protein expression by 1 mM isopropyl β-D-thiogalactopyranoside. After 4h of induction, cultures were centrifuged at 6000 x g, for 10 min, at 20°C and the pellets were resuspended in lysis buffer (50 mL *per* liter of bacterial culture) containing 20 mM HEPES, 300 mM NaCl, 6 M urea pH 7.2 before being frozen at -20°C. The suspensions were then thawed and sonicated for 5 min at 600 watts and the lysates were then centrifuged for 20 min at 15 000 x g at 10°C.
In the case of the W proteins and HeV Cys variants, the supernatants were mixed with Ni-NTA beads (2.5 mL of beads *per* 50 mL of lysate, Cytiva) pre-equilibrated in lysis buffer, and incubated under gentle agitation for 1 h at room temperature (RT). The resin was then washed with lysis buffer supplemented with 1 M NaCl and 10 mM imidazole, and then re-equilibrated with desalting buffer containing 20 mM HEPES, 50 mM NaCl, 6 M urea, 10 mM imidazole pH 7.2. Proteins were eluted with desalting buffer supplemented with 250 mM imidazole. Recombinant proteins were then reduced by addition of 10 mM DTT followed by incubation for 30 min at RT. Recombinant proteins were further purified by anion exchange chromatography (AEC) on a HiPrep DEAE FF anion exchange column (Cytiva) equilibrated in desalting buffer without imidazole. The proteins were eluted with an NaCl gradient (50 – 500 mM) in desalting buffer. All the recombinant proteins were eluted at around 100 mM NaCl. Purified samples obtained from (AEC) were concentrated by centrifugation with an amicon ultra-15 with a 10 kDa cut-off (Millipore) before being reduced for 30 min by 5 mM DTT at RT. DTT was then removed by injecting proteins samples on a HiPrep 26/10 desalting column (Cytiva) equilibrated in 20 mM HEPES, 150 mM NaCl, 6 M urea pH 7.2 (HBS/6 M urea buffer, storage buffer). Purified proteins were directly aliquoted and stored at -80°C.
FKBP- $C^{all}S$ was purified in the same way as the W proteins and $W^{HeV}$ Cys variants except that DTT was omitted, and that an additional preparative SEC step was performed after the AEC. Since the FKBP moiety contains a single Cys residue, FKBP- $C^{all}S$ was reduced by 10 mM DTT for 30 min at RT before being injected on a HiLoad 16/600 Superose 6 pg column equilibrated in HBS/1 M urea, pH 7.2. Purification was carried out at a flow rate of 1 mL/min, at room temperature. Eluted samples were concentrated as described above, aliquoted and stored at -80°C.
Except for the Ni-NTA affinity chromatography, all the purification steps were carried out on an ÄKTA Pure Protein Purification System (Cytiva) operated at room temperature. Purified proteins analyzed by SDS-PAGE are shown in **Supplementary Figure S9**. Protein concentration was determined spectrophotometrically at 280 nm using the molar extinction coefficient calculated with ProtParam. Protein

amino acid numbering is according to the UniProt database ($W^{HeV}$ and $W^{NiV}$ accession numbers P0C1C6 and P0C1C7, respectively).

**Protein sequencing by tandem mass spectrometry**

The identity of the compact conformation of $W^{HeV}$ (*i.e.* peaks II in **Supplementary Figure S2B**) was confirmed by mass spectrometry analysis of tryptic fragments obtained after digestion of the purified protein band excised from SDS-polyacrylamide gel. The excised band was analyzed by the mass spectrometry facility of Marseille Proteomics. Briefly, gel bands were cut, destained in 100 mM $NH_4HCO_3$ in 50 % acetonitrile, dried at room temperature, then rehydrated. Cysteines were then reduced using 10 mM DTT in 100 mM $NH_4HCO_3$ pH 8.0 for 45 min at 56°C before alkylation in the presence of 55 mM iodoacetamide in 100 mM ammonium bicarbonate pH 8.0 for 30 min at room temperature in the dark, and bands were finally washed twice with 25 mM $NH_4HCO_3$ pH 8.0 and digested with high-sequencing-grade trypsin (Promega, Madison, WI). Mass spectrometry analyses were carried out by LC-MSMS using a Q Exactive Plus Hybrid Quadrupole-Orbitrap online with a nanoLC Ultimate 3000 chromatography system (Thermo Fisher Scientific™, San Jose, CA). For each sample, 3 microliters were injected on the system. After pre-concentration and washing of the sample on a Acclaim PepMap 100 column (C18, 2 cm × 100 µm i.d. 100 A pore size, 5 µm particle size), peptides were separated on a LC EASY-Spray column (C18, 50 cm × 75 µm i.d., 100 A, 2 µm, 100A particle size) at a flow rate of 300 nL/min with a two steps linear gradient (2-22% acetonitrile/$H_2O$; 0.1 % formic acid for 100 min and 22-32% acetonitrile/$H_2O$; 0.1 % formic acid for 20 min). For peptide ionization in the EASYSpray source, spray voltage was set at 1.9 kV and the capillary temperature at 250 °C. All samples were measured in a data dependent acquisition mode. Each run was preceded by a blank MS run in order to monitor system background. The peptide masses were measured in a survey full scan (scan range 375-1500 m/z, with 70 K FWHM resolution at m/z=400, target AGC value of 3.00×106 and maximum injection time of 100 ms). Following the high-resolution full scan in the Orbitrap, the 10 most intense data-dependent precursor ions were successively fragmented in HCD cell and measured in Orbitrap (normalized collision energy of 25 %, activation time of 10 ms, target AGC value of 1.00×103, intensity threshold 1.00×104 maximum injection time 100 ms, isolation window 2 m/z, 17.5 K FWHM resolution, scan range 200 to 2000 m/z). Dynamic exclusion was implemented with a repeat count of 1 and exclusion duration of 20 s. Raw files generated from mass spectrometry analysis were processed with Proteome Discoverer 1.4 .1.14 (Thermo Fisher Scientific, San Jose, CA) to search against a homemade database containing 20150 human sequences, 4306 *E. coli* sequences implemented with the eight expected sequences. (swissprot – human – reviewed – 170315 _ 20150 _UP_coli_171120_4306_ Patrick220922 _ ID _ Bandes.fasta). Database search with SequestHT were done using the following settings: a maximum of two trypsin miscleavage allowed, methionine oxidation and N terminal protein acetylation as variable modifications, and cysteine carbamidomethylation as fixed modification. A peptide mass tolerance of 6 ppm and a fragment mass tolerance of 0.8 Da were allowed for search analysis. Only peptides with high Sequest scores were selected for protein identification. False discovery rate was set to 1 % for protein identification. The selected peptides were then compared to the theoretical digestion peptides of the database leading to the identification of the major protein of the band by generating a Sequest score.

**Turbidimetry and fluorimetry measurements**

$W^{HeV}$ (WT and variants) and $W^{NiV}$ (WT) protein samples of 500 µL at a concentration of 75 µM in HBS/6 M urea buffer were incubated at 37°C for different times (indicated in each experiment). Protein solutions were then buffer exchanged to remove urea using a PD Minitrap G-25 column (Cytiva) equilibrated in HBS only. The resulting urea-free protein samples (1000 µL) were analyzed at a final concentration of 30 µM in the presence of 50 µM thioflavin T (ThT). Turbidity measurements, made at 360 nm, and fluorescence measurements were performed on a Tecan microplate reader GENios Plus after 5 min of

temperature equilibration at 37°C. Measurements were then made every 30 min for 120 min in black 96-well plates with transparent flat bottom (Greiner, 655096) incubated at 37°C between each measurement. ThT was excited at 430 nm and fluorescence emission was recorded at 535 nm.
The same protocol was followed in the experiments made in the presence of DTT. 2 mM DTT was added either just after the buffer exchange step ($t_0$), or on a $W^{HeV}$ sample that had already been incubated for two hours ($t_{2h}$). In this latter case, turbidity and fluorescence were monitored for two extra hours ($t_{2h + 2h}$). This latter sample was further incubated at 37°C for a total of 24h and then centrifuged at 10 000 x g for 10 min. The pellet was resuspended in 150 µL of HBS before being deposited on a NS-EM grid.
Measurements were performed at least in triplicate (indicated in each experiment) and data are represented as mean ± standard deviation (s.d.). The statistical analysis used is detailed for each experiment, using either a one-way ANOVA with the Dunnett's multiple comparison test, or a Student's t-test.

**Negative-staining electron microscopy (NS-EM)**

Proteins incubated for 120 min at 37°C in HBS for aggregation and ThT binding measurements were analyzed by NS-EM. For this purpose, protein samples were homogenized by pipetting and diluted in HBS at a final concentration of 2 µM. Carbon coated grids (Carbon 300 mesh 3mm Cu, TAAB) were exposed to plasma glow discharge for 20 seconds using a GloQube (Quorum) with a current of 25 mA in order to increase protein adhesion. Three µL of sample were deposited on the grid for a minute, then blotted and washed three times in HBS before being incubated with 35 µL of 1 % (w/v) uranyl acetate solution for one minute. Grids were then blotted and dried overnight before being observed on a TECNAI T12 Spirit microscope (Thermo company) operated at 120 kV equipped with a Veleta 2Kx2K CCD camera (Olympus). Measurements of fibril lengths were made manually with the ImageJ software (Schneider et al., 2012).

**Chemically-induced dimerization of FKBP-$C^{all}$S**

266 µL of a FKBP-$C^{all}$S sample were incubated for 10 min at a final concentration of 70 µM in HBS containing 400 mM urea at RT in the absence or presence of 140 µM AP20187 (Sigma-Aldrich, SML2838). After 10 min, 100 µL of undiluted sample were injected onto a Superdex S200 Increase 10/300 GL column (Cytiva) at a flow rate of 0.8 mL/min at RT, for analytical purposes. In parallel, the rest of the sample was buffer exchanged using a PD Minitrap G-25 column (Cytiva) equilibrated in HBS to remove excess AP20187. Buffer exchanged samples were analyzed by turbidity and fluorimetry in the presence of ThT at a final concentration of 30 µM in the same way as described above. After 2h of incubation, NS-EM grids were prepared at a final concentration of 2 µM in HBS as described above.

**$W^{HeV}$ $C^{all}$S variant crosslinking by DTSSP**

720 µL of $W^{HeV}$ $C^{all}$S at a final concentration of 110 µM in HBS/6 M urea were cross-linked in the presence of 2 mM 3,3'-dithiobis(sulfosuccinimidyl propionate), DTSSP (Pierce, 21578) for 40 min at room temperature. The DTSSP contains a primary amine-reactive group at each end and a 8-carbon spacer arm. The reaction was stopped by addition of 100 µL of a 1 M Tris HCl pH 8.0 solution for 15 min at room-temperature. The resulting sample was analyzed by SEC at 75 µM, as well as by turbidimetry and fluorimetry after a buffer exchange step in HBS, as described above, at a final concentration of 30 µM. NS-EM grids (with samples at 2 µM) were prepared after 2h of incubation in HBS.

**Immunofluorescence**

For W immunofluorescence of ReAsH-stained cells, HEK293T cells were seeded in Ibidi µ-Slide 8 Well (Ibidi, Cat# 80826) previously coated with 50 µg/mL poly-D-lysine (Sigma-Aldrich, Cat# P6407). Cells were transfected 24h later with 0.4 µg of an empty pcDNA3.1(+) vector, a pIRES2-EGFP plasmid driving Enhanced green fluorescent protein (EGFP) expression, or pcDNA3.1(+) plasmids coding for WT $W^{HeV}$ fused to a tc tag either at the N- or C-terminus, using the TransIT-LT1 reagent (Mirus, Cat# MIR 2300) with a DNA:transfection reagent ratio of 1:3 according to the manufacturer's instructions. The TC-ReAsH™ II In-Cell Tetracysteine Tag Detection Kit (Invitrogen, Cat# T34562) was then used to stain the tc tag at 24h post-transfection, according to the manufacturer's instructions. The ReAsH-EDT2 labeling reagent was used at a final concentration of 1.25 µM. Cells were then fixed with 4% methanol-free formaldehyde (Electron Microscopy Sciences, Cat# 15713) for 10 min, washed with Dulbecco's Phosphate Buffered Saline (DPBS) (Gibco, Cat# 14190-144), and permeabilized with 0.1% Triton X-100 (Sigma-Aldrich, Cat# T8787)-3% bovine serum albumin (BSA) (Sigma-Aldrich, Cat# A3059) in DPBS for 15 min. Next, cells were incubated for 1 h at room temperature with a rabbit anti-$W^{CTD}$ antibody (GeneScript) (1:100) in DPBS-0.1% Triton X-100-3% BSA, rinsed thrice with DPBS for 3 min and incubated for 1 h at room temperature with Alexa Fluor 647 Donkey anti-Rabbit IgG (H+L) Highly Cross-Adsorbed Secondary antibody (Invitrogen, Cat# A-31573) (1:500) in DPBS-0.1% Triton X-100-3% BSA. Following three washes of 3 min with DPBS, cells were incubated with 1 µg/mL 4',6-diamidino-2-phenylindole (DAPI) for 5 min and washed thrice with DPBS before imaging on a Yokogawa HCS CQ1 confocal system (PLATIM, SFR Biosciences Lyon). Z-stacks of 10-µm-range with an imaging step of 0.8 µm were acquired with the 40X objective. Maximum intensity projections were generated automatically by the microscope software. A similar protocol was used for Figure S5, except cells were not stained with ReAsH.

**Embedding and ultramicrotomy of transfected cells, and EM analyses**

HEK293T cells were washed 4 times in PBS and 4 times in $H_2O$ for 5 minutes each. For contrast and fixation, the sample was incubated with 1% osmium tetroxide ($OsO_4$) in $H_2O$ for 1h at room temperature. The samples were then rinsed 4 times in $H_2O$. An additional fixation and contrast step was performed in 1 % uranyl acetate in $H_2O$ for 1h at 4°C, and the samples were rinsed 3 times in $H_2O$. For sample dehydration, cells were incubated with increasing concentrations of ethanol (25%, 50%, 70%, 80%, 90%, 100%) for 10 minutes each, plus a final incubation in 100 % ethanol for 20 minutes. For resin infiltration, cells were placed in 1/4 Epon 812 LX112, 3/4 ethanol overnight at 4°C. The next day, the HEK293T cells were first incubated in 1/2 Epon 812 LX112, 1/2 ethanol and then in 3/4 Epon 812 LX112, 1/4 ethanol (1h each, room temperature). Finally, the cells were incubated in pure Epon 812 LX112 overnight. The next day, the pure Epon 812 LX112 was replaced with fresh one and incubated for 4h at room temperature. Lastly, the resin was replaced with fresh Epon 812 LX112 and incubated at 60°C for approximately 24h, allowing the polymerization to take place.

**Supplementary Table S1.** Length distribution of fibrils formed by $W^{HeV}$. Proteins were preincubated in oxidative conditions for various times at 37°C in urea before urea removal and then incubated for 2h in urea-free buffer.

| $W^{HeV}$ & variants | Number of fibrils analyzed | Mean length (nm) | Median (nm) | Minimum length (nm) | Maximum length (nm) |
|---|---|---|---|---|---|
| $W^{HeV}$ $t_{24}$ | 193 | 273 | 249 | 50 | 834 |
| $W^{HeV}$ $t_{48}$ | 306 | 255 | 245 | 36 | 661 |
| $W^{HeV}$ $t_{72}$ | 330 | 179 | 170 | 7 | 620 |

**Supplementary Table S2.** Percentage of cells with filaments as a function of *tc* tag position.

| $W^{HeV}$ variant | Cells with filaments (%) 24h pt | Cells with filaments (%) 48h pt |
|---|---|---|
| *tc*-$W^{HeV}$ | ~6% | ~6% |
| $W^{HeV}$-*tc* | ~3% | ~3% |

**Supplementary Table S3.** Templates and primers used for the generation of DNA constructs.

| Variant | Template[a] for PCR 1 and 2 | Primers for PCR1 | Primers for PCR 2 |
|---|---|---|---|
| **C123$^{only}$** | HWC316S (and HWC419S) | attb1 + HWC316SRlong | HWC334Sflong + attb2 |
| **C316$^{only}$** | HWC123S (and HWC419S) | attb1 + HWC316SR | HWC316SF + attb2 |
| **C334$^{only}$** | C334C419 | attb1 + HWC419SR | HWC419SF + attb2 |
| **C419$^{only}$** | HWC123S | attb1 + HWC316SRlong | HWC334Sflong + attb2 |
| **W$^{HeV}$C$^{all}$S** | HWC123S[c] (and HWC419S[d]) | attb1 + HWC316SRlong | HWC334Sflong + attb2 |
| **W$^{HeV}$Δ29** | HW | HPNT1d29B1TEV | attb2 |
| **W$^{HeV}$Δ29C$^{all}$S** | HWCallS | HPNT1d29B1TEV | attb2 |
| **FKBP-C$^{all}$S** | FKBP (and C$^{all}$S) | FKBPb1 + FKBP3 | FKBP2 + attb2 |
| **W-*tc*** **(in pcDNA3.1(+)** | HW | HindHP + HW4CXho | |
| ***tc*-W** **(in pcDNA3.1(+)** | HW | HW4CHind + XhoHW | |
| ***tc*-W** | pCG-FLAG-W (unpublished) | HeVtc-W F + HeVtc-W R | |
| ***tc*-W$^{CallS}$** | N/A, DNA fragment was synthetized by ThermoFisher Scientific GeneArt service | | |
| ***tc*-WΔ29** | pCG-FLAG-W (unpublished) | HeVtc-Wd29 F + HeVtc-W | |
| ***tc*-WΔ29$^{CallS}$** | N/A, DNA fragment was synthetized by ThermoFisher Scientific GeneArt service | | |
| ***tc*-mStayGold** | N/A, DNA fragment was synthetized by ThermoFisher Scientific GeneArt service | | |

[a]Except for FKBP and eukaryotic constructs, all DNA templates are borne by the Gateway vector pDEST17O/I.

[b]used in PCR 1.

[c]used in PCR 2.

FKPB Uniprot accession number: P62942 with a F37V substitution. No linker between the FKBP and W$^{HeV}$ C$^{all}$S moieties.

**Supplementary Table S4.** Sequence of the primers used to generate the constructs used in this work.

| Primer | Sequence |
|---|---|
| **attb1** | ACAAGTTTGTACAAAAAAGCAGGCT |
| **attb2** | ACCACTTTGTACAAGAAAGCTGGGT |
| **HWC123SF** | GTATCATGATCATGGTGGCGAAAGCACCGGCCATGGCCCGAGCAG |
| **HWC123SR** | TTCGCCACCATGATCATGATAC |
| **HWC316SF** | GATAGCCTGATGCAGGATAGCAGTAAACGTGGTGGCGTGCCG |
| **HWC316SR** | AGGTTTTGCGGCCCAGCATG |
| **HWC419SF** | CATGCTGGGCCGCAAAACCAGCCTGGGCCGCCGCGTGGTGC |
| **HWC419SR** | TGGTTTTGCGGCCCAGCATG |
| **HWC334Sflong** | GAAACGTCTGCCGATGCTGAGCGAAGAATTCGAATCGAGCGGTAGCGATGATCCG |
| **HWC316SRlong** | TCAGCATCGGCAGACGTTTCGGCACGCCACCACGTTTACTGCTATCCTGCATCAGGC |
| **HWC316SF** | GCTGAGCGAAGAATTCGAAAGCAGCGGTAGCGATGATCCGATT |
| **HWC316SR** | TTCGAATTCTTCGCTCAGC |
| **FKBPb1** | ACAAGT TTGTACAAAAAAGCAGGCTCGGGAGTGCAGGTGGAGACTATC |
| **FKBP2** | GTGGAGCTTCTAAAACTGGAAGATAAACTGGATCTGGTTAAC |
| **FKBP3** | TTCCAGTTTTAGAAGCTCCAC |
| **HPNT1d29B1TEV** | ACAAGTTTGTACAAAAAAGCAGGCTCTGAAAACCTGTACTTCCAGGGTCGTAGCAGCATTCAGCAGCCGAGC |
| **HindHP** | GCGTTTAAACTTAAGCTTCCACC ATG GATAAACTGGATCTGGTTAACG |
| **HW4CXho** | GGGCCCTCTAGACTCGAGTTATTAGGGCTCCATGCAGCAGCCGGGGCAGCAGTTCAGGAAGTTGCTCATGCGGCGCAGCAGCAC |
| **HW4CHind** | GTTTAAACTTAAGCTTCCACCATGTTCCTGAACTGCTGCCCCGGCTGCTGCATGGAGCCCGATAAACTGGATCTGGTTAACG |
| **XhoHW** | GCCCTCTAGACTCGAGTTATTAGTTGCTCATGCGGCG |
| **HeVtc-W F** | CAAAGAATTCCTCGAGCGGCCGCCACCATGTTCCTGAACTGCTGCCCCGGCTGCTGCATGGAGCCCGACAAGTTGGATCTAGTCAA |
| **HeVtc-Wd29 F** | CAAAGAATTCCTCGAGCGGCCGCCACCATGTTCCTGAACTGCTGCCCCGGCTGCTGCATGGAGCCCCGATCAAGCATCCAACAACC |
| **HeVtc-W R** | GAGTGAATTCCTCGAGACGCGTCAGTTGGACATTCTCCGCA |

### SUPPLEMENTARY FIGURES

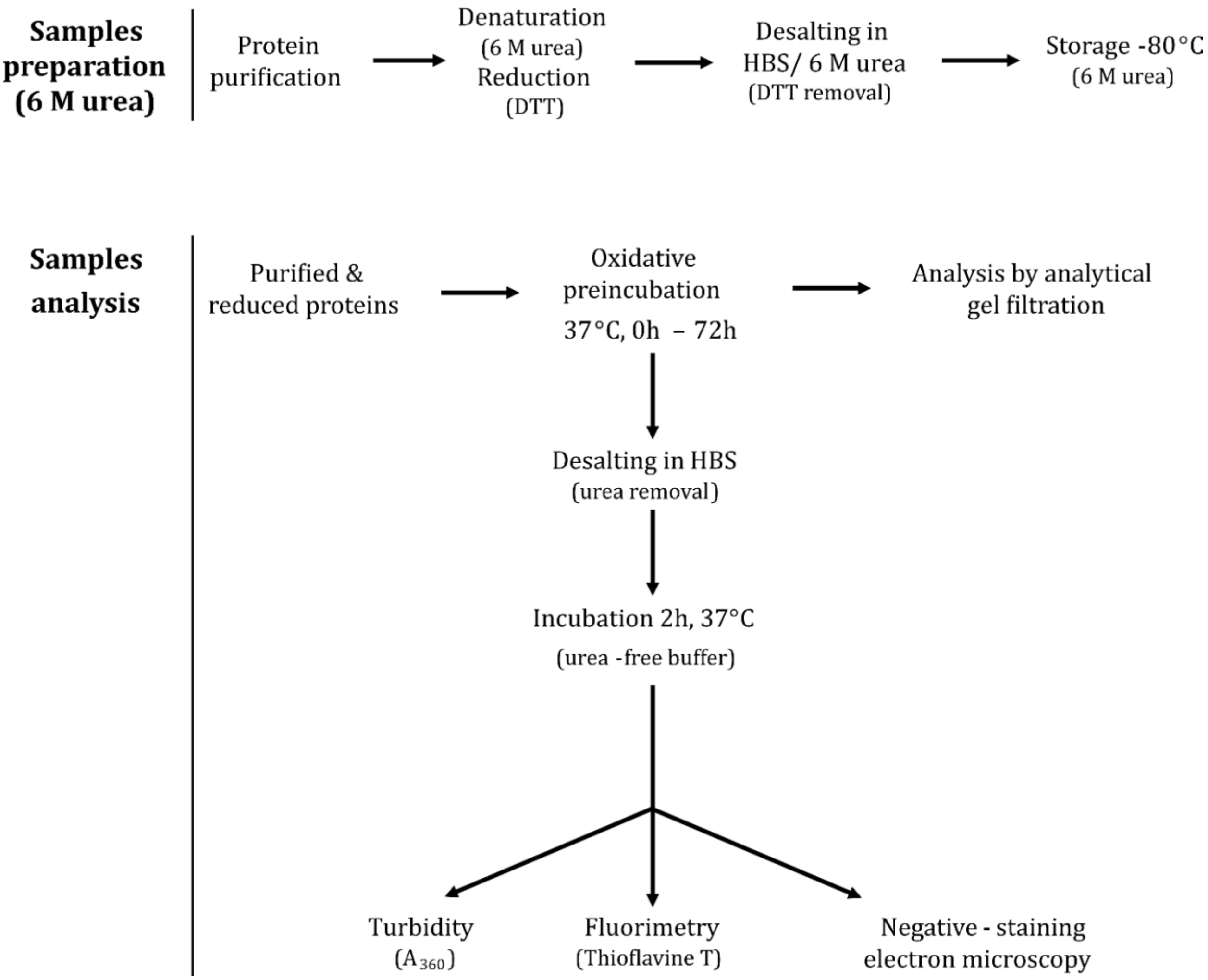


**Supplementary Figure S1**. **Experimental procedures regarding the preparation and the analysis of recombinant proteins used in this work.** All proteins were purified in denaturing conditions and reduced to recover only monomeric species and to abrogate their aggregation. Natural oxidation was carried out in urea and the denaturing buffer (HBS/6 M urea) was exchanged for an urea-free buffer (HBS) to induce protein aggregation, that was followed for 2h by turbidimetry, thioflavin T (ThT) fluorescence and negative-staining electron microscopy (NS-EM).

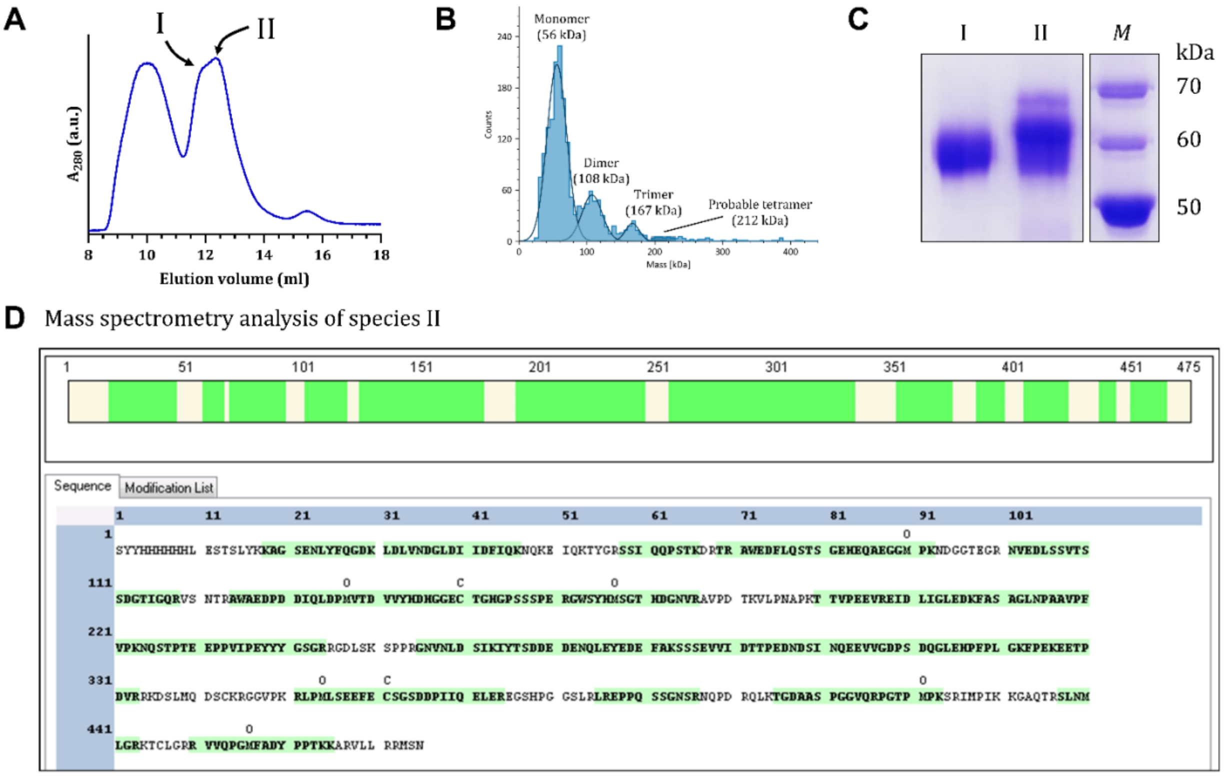


**Supplementary Figure S2. Identification of W^HeV compact conformations and of its oligomers by mass spectrometry.** Analytical SEC (**A**) and mass photometry **(B)** analysis of W^HeV preincubated for 48h in urea. Peak I corresponds to the initial monomeric species. Species II (whose formation is cysteine-dependent) is eluted at ≈ 12.3 mL. Dimeric species and oligomers are eluted around 10 and 9-9.5 mL, respectively. (**C)** Non-reducing SDS-PAGE analysis of species I and II. (**D)** Proteolytic peptides identified by mass spectrometry analysis of species II (leading to a coverage of 71.9 %) of the W^HeV sequence.

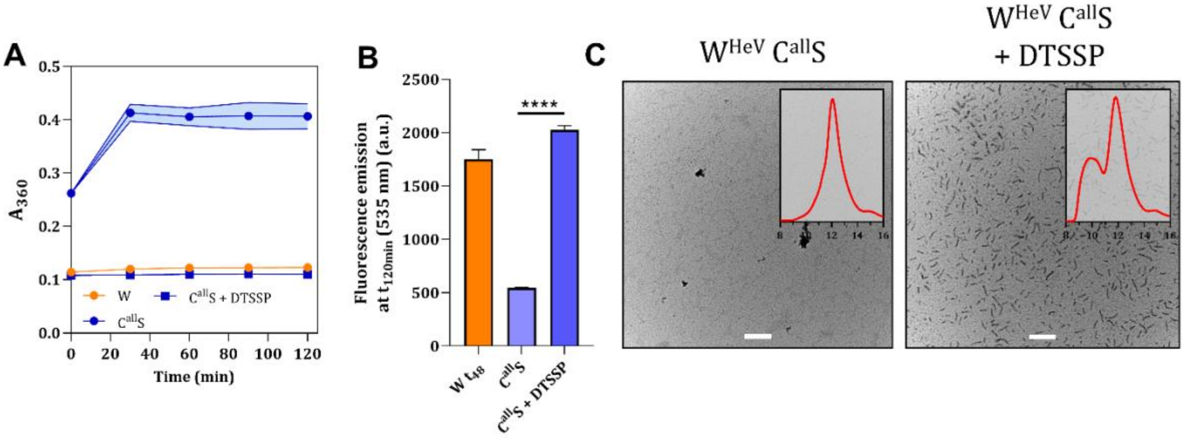


**Supplementary Figure S3. Analysis of $W^{HeV}$ $C^{all}S$ reticulated with DTSSP.** Turbidimetry (**A)** and ThT fluorescence measurements (**B)** of non-cross-linked (blue circles), and cross-linked (blue squares) $W^{HeV}$ $C^{all}S$. WT $W^{HeV}$ data (W, orange circles) preincubated for 48h are shown for comparison. (**C)** NS-EM micrographs of $W^{HeV}$ $C^{all}S$ with or without cross-linking. The insets show the corresponding analytical SEC profiles. Panels A and B show mean values and s.d. as obtained from n=4 independent measurements. The shaded region in (B) corresponds to the error bar. Statistical analysis was made with a Student's t-test; ****: $p<0.0001$. Scale bar: 200 nm.

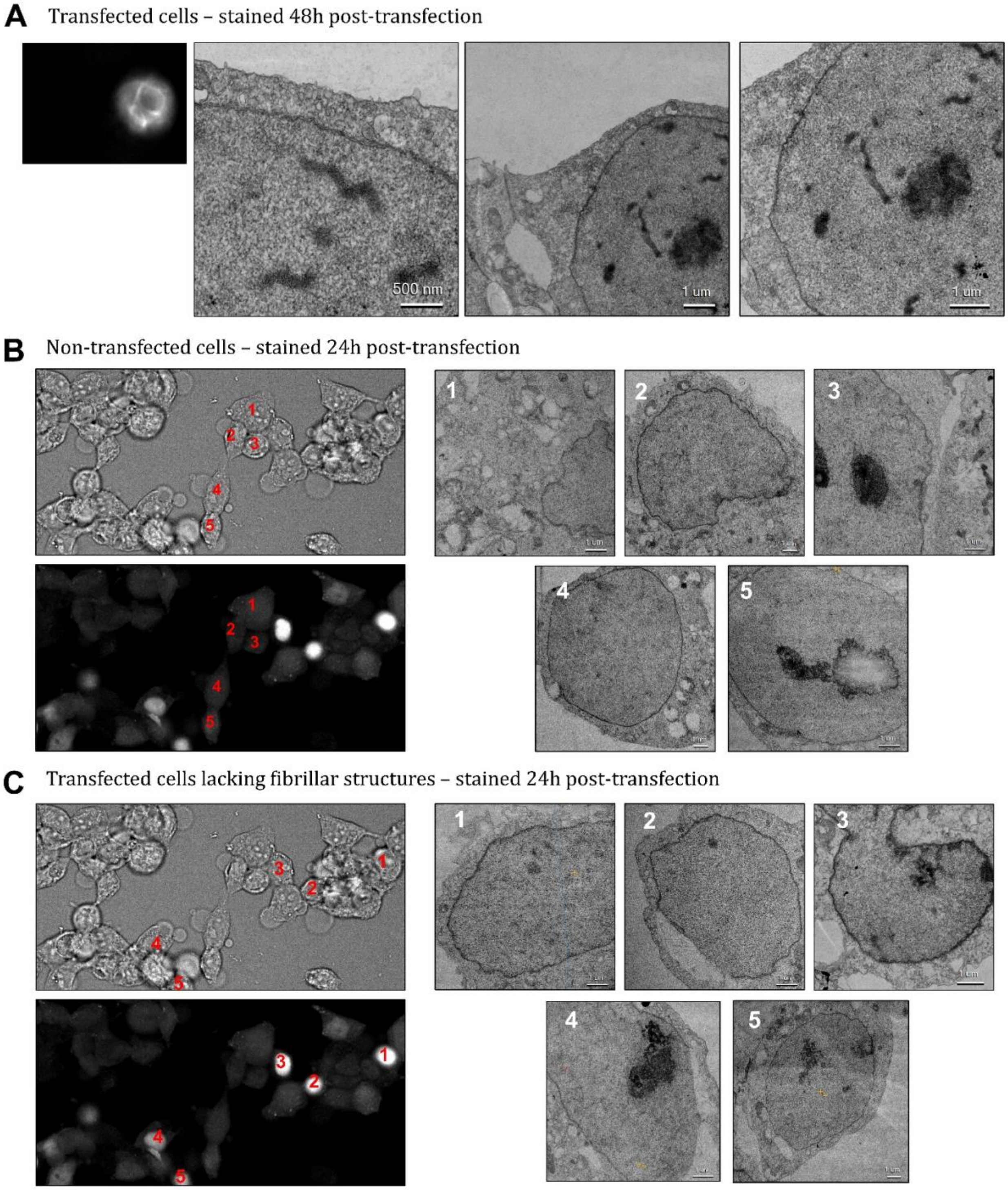


**Supplementary Figure S4. Z-stack projections of HEK293T cells transfected with tetracysteine (*tc*)-tagged $W^{HeV}$ constructs with relative ultrathin sections EM images.** Cells were stained 24h and 48h post-transfection with ReAsH dye to detect the *tc* peptide and imaged by confocal microscopy. (**A**) Z-stack slice of HEK293T cells stained 48h post transfection and electron micrograph of the corresponding ultrathin section. (**B**) Z-stack slice of HEK293T cells stained 24h post transfection and the electron micrograph of the corresponding ultrathin section. Five control cells (*i.e.* lacking signal for the tetracysteine-labelled $W^{HeV}$ constructs), selected from a pool of 40 visualized cells, are shown (see numbers in red). (**C)** Z-stack slice of HEK293T cells stained 24h post transfection and the electron micrograph of the corresponding ultrathin section. Five cells positive for transfection (but lacking fibrillar structures visible in the Z-stack projection) selected from a pool of 35 visualized cells are shown (see numbers in red).

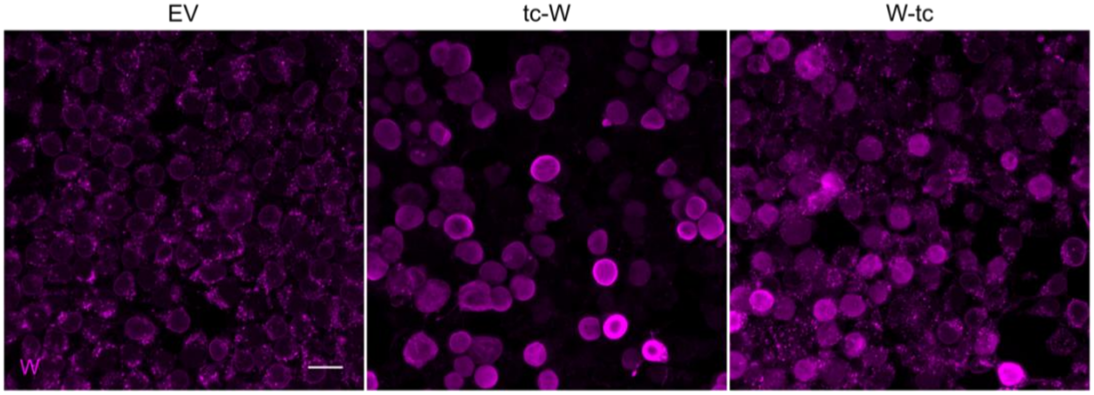


**Supplementary Figure S5. IF analysis of $W^{HeV}$ expression in transfected HEK293T cells.** The W expression level in HEK293T cells transfected with the indicated constructs was analyzed by IF staining (anti-W antibody) and confocal microscopy at 24h post-transfection as described in Figure 4A. EV: empty vector. Scale bar, 20 µm.

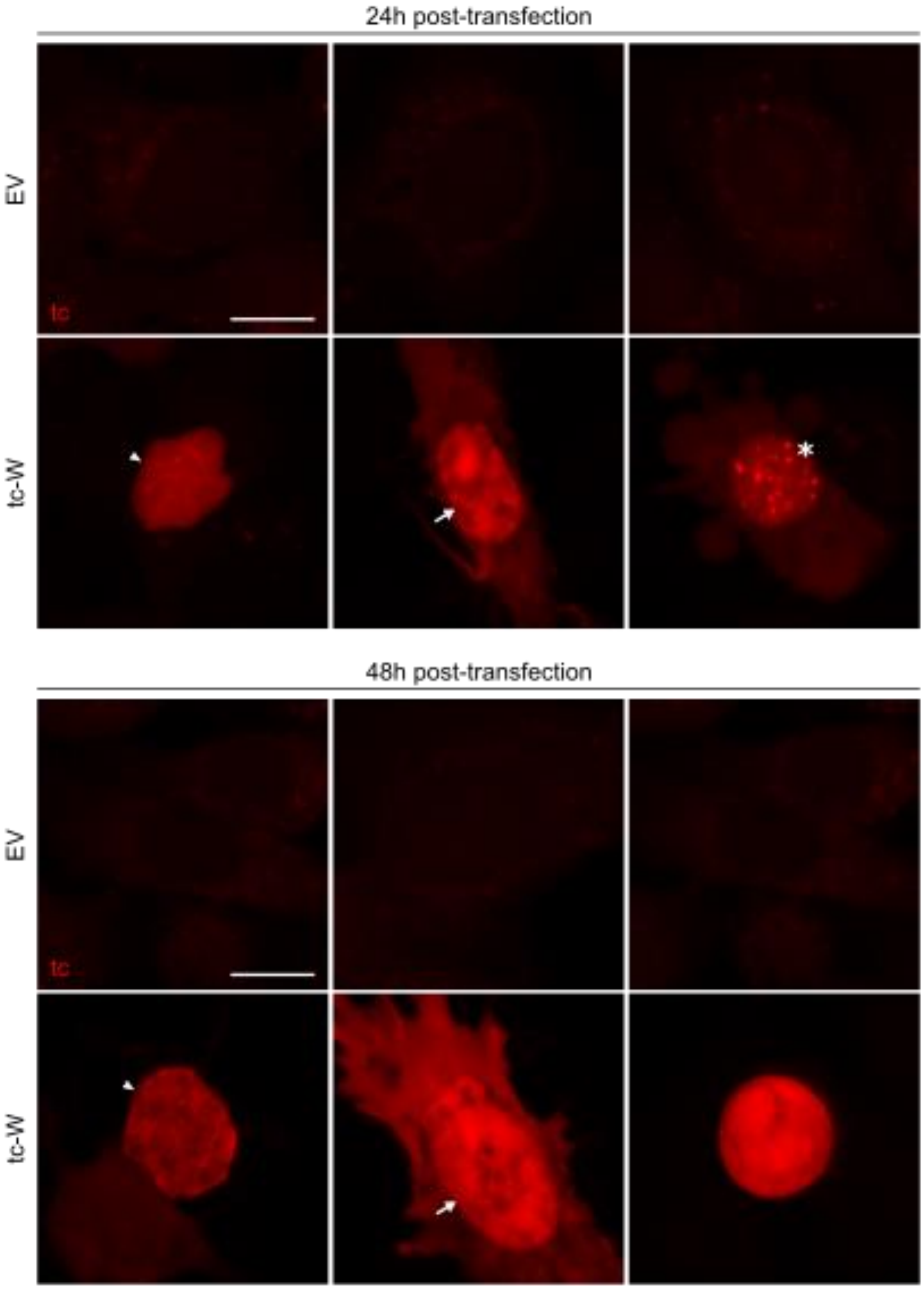


**Supplementary Figure S6.** W forms nuclear filamentous structures and other types of condensates in A549 cells. Z-stack projections of A549 cells transfected with an EV or the *tc*-W construct. Cells were stained 24h and 48h post-transfection with ReAsH dye and imaged by confocal microscopy. Note the presence of filaments (arrowheads) along with aggregates (arrows) and spots (asterisk). Scale bar, 10 µm.

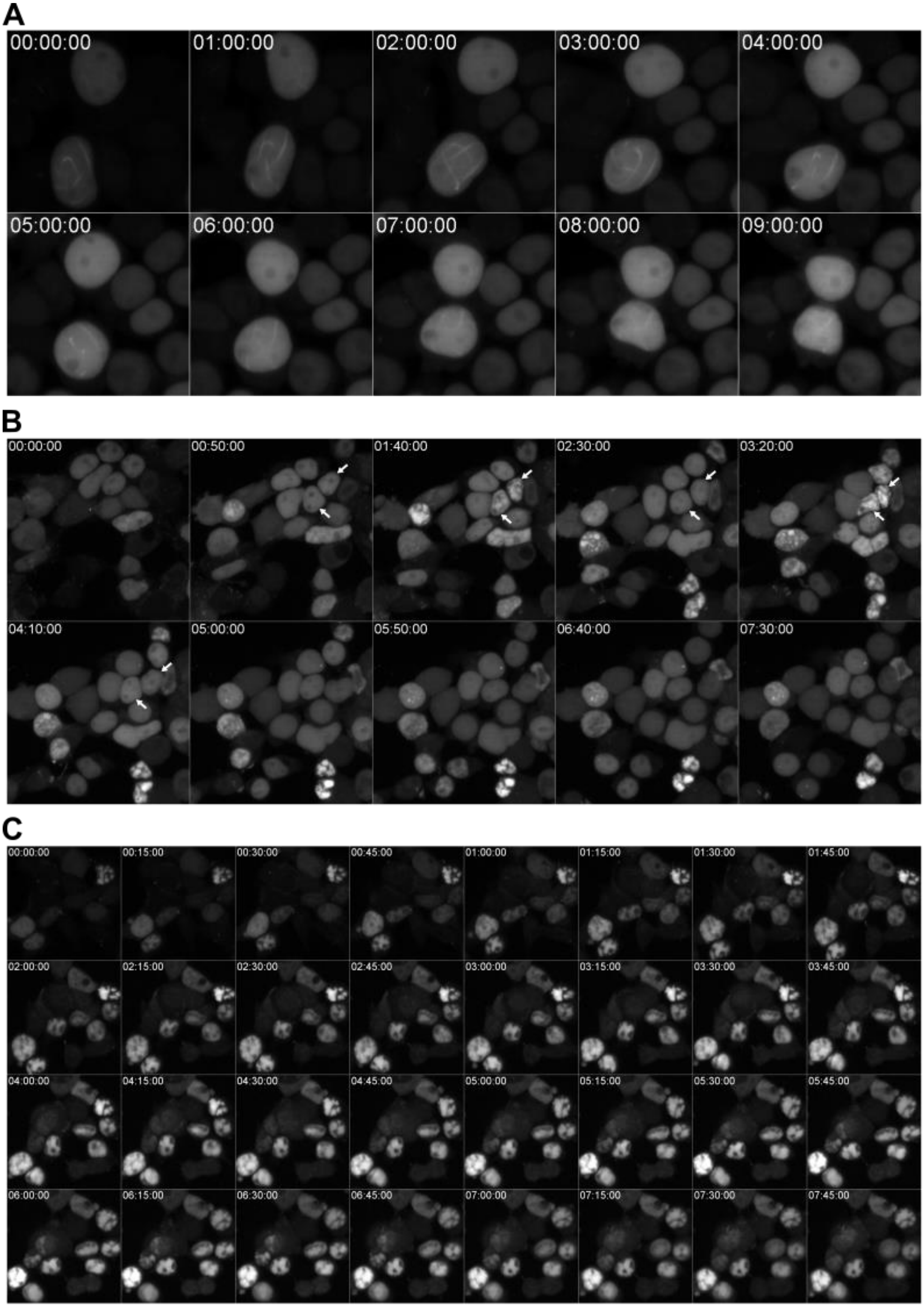
A
00:00:00
01:00:00
02:00:00
03:00:00
04:00:00
05:00:00
06:00:00
07:00:00
08:00:00
09:00:00
B
00:00:00
00:50:00
01:40:00
02:30:00
03:20:00
04:10:00
05:00:00
05:50:00
06:40:00
07:30:00
C
00:00:00
00:15:00
00:30:00
00:45:00
01:00:00
01:15:00
01:30:00
01:45:00
02:00:00
02:15:00
02:30:00
02:45:00
03:00:00
03:15:00
03:30:00
03:45:00
04:00:00
04:15:00
04:30:00
04:45:00
05:00:00
05:15:00
05:30:00
05:45:00
06:00:00
06:15:00
06:30:00
06:45:00
07:00:00
07:15:00
07:30:00
07:45:00

**Supplementary Figure S7.** Time-lapse imaging of transfected HEK293T cells showing the dynamics of $W^{HeV}$ filaments and aggregates. (**A-C)** Z-stack projections of live HEK293T cells transfected with *tc*-$W^{HeV}$ construct, and imaged by confocal microscopy in the presence of FlAsH, at the indicated times (in the format h:min:sec). (**A**) Evolution of the morphology of a nuclear filament, which seems to progressively disappear. (**B**) Dynamics of the reversible amorphous condensation of W, as shown with nuclei alternately harboring a diffuse homogeneous signal and amorphous aggregates (arrows). (**C)** Morphology of multiple nuclei displaying amorphous aggregates with different aspects.

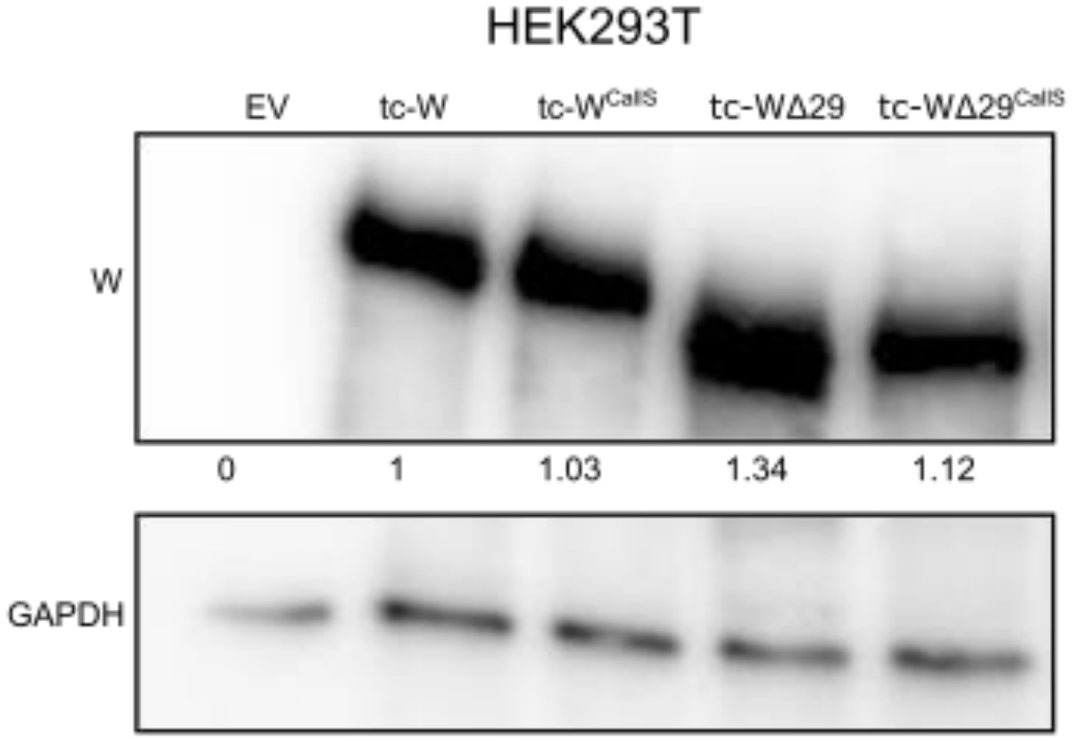


**Supplementary Figure S8.** Western blot analysis of the expression of the $W^{HeV}$ variants in transfected HEK293T cells at 24h post-transfection.

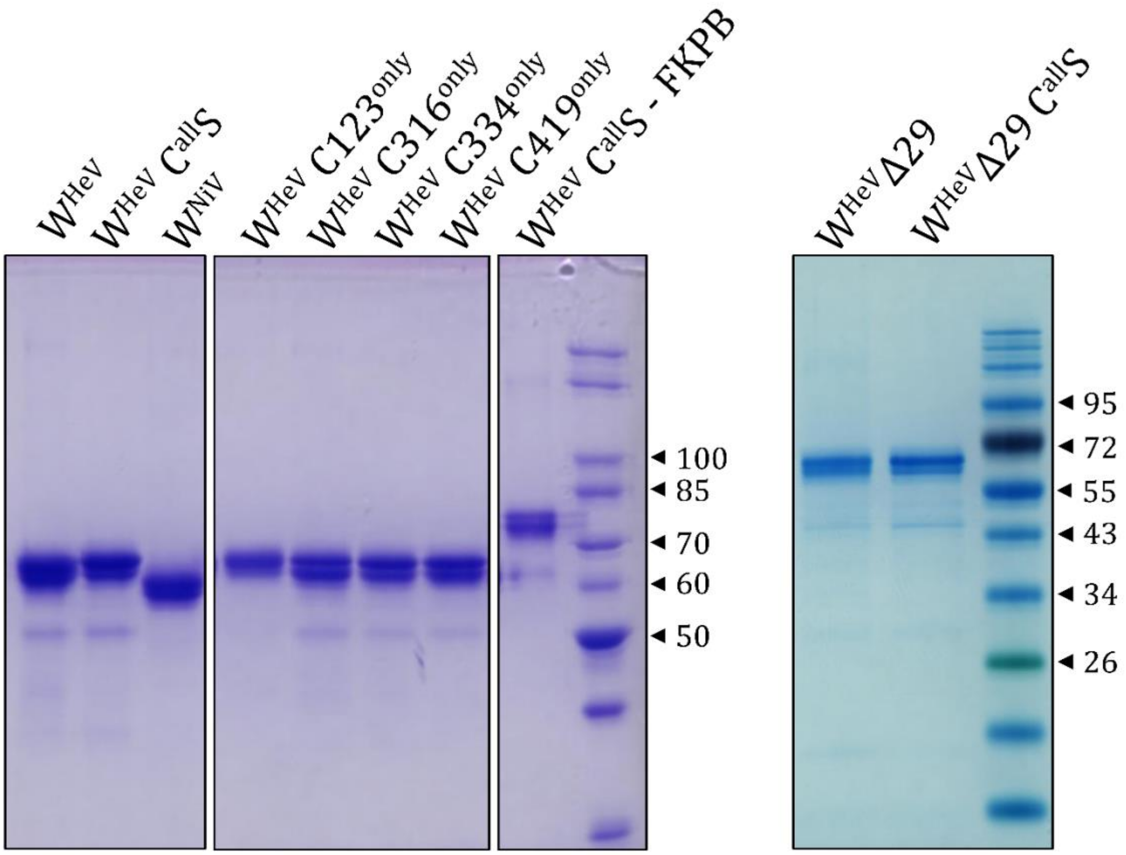


**Supplementary Figure S9. SDS-PAGE analysis of all the recombinant proteins used in this work.** 2 µg of the W proteins and of their variants were analyzed under reducing conditions (5 mM DTT) on a 10 % polyacrylamide gel. Gels were stained with Coomassie Brillant Blue R-250. Molecular markers: New England Biolabs, P7717S (left), P7719S (right).

**Supplementary Movie S1:** 3D visualization of $W^{HeV}$ filaments in nuclei of transfected HEK293T cells. Cells transfected with the *tc*-W construct were imaged by confocal microscopy 72h post-transfection as previously described, and visualized using Fiji.

**Supplementary Movie S2:** 3D visualization of $W^{HeV}$ globular condensates in nuclei of transfected HEK293T cells. Cells transfected with the *tc*-W construct were imaged by confocal microscopy 72h post-transfection as previously described, and visualized using Fiji.
